# A Population Representation of the Confidence in a Decision in the Parietal Cortex

**DOI:** 10.1101/2024.08.15.608159

**Authors:** Ariel Zylberberg, Michael N. Shadlen

**Affiliations:** Mortimer B Zuckerman Mind Brain Behavior Institute, Columbia University, New York, United States; Virtual Confidence and Metacognition Laboratory; Department of Neuroscience, Columbia University, New York, United States; The Kavli Institute for Brain Science, Columbia University, New York, United States; Howard Hughes Medical Institute, Chevy Chase, United States

## Abstract

Confidence in a decision is the belief, prior to feedback, that one’s choice is correct. In the brain, many decisions are implemented as a race between competing evidence-accumulation processes. We ask whether the neurons that represent evidence accumulation also carry information about whether the choice is correct (i.e., confidence). Monkeys performed a reaction time version of the random dot motion task. Neuropixels probes were used to record from neurons in the lateral intraparietal (LIP) area. LIP neurons with response fields that overlap the choice-target contralateral to the recording site (T_in_ neurons) represent the accumulation of evidence in favor of contralateral target selection. We demonstrate that shortly before a contralateral choice is reported, the population of T_in_ neurons contains information about the accuracy of the choice (i.e., whether the choice is correct or incorrect). This finding is unexpected because, on average, T_in_ neurons exhibit a level of activity before the report that is independent of reaction time and evidence strength—both strong predictors of accuracy. This apparent contradiction is resolved by examining the variability in neuronal responses across the population of T_in_ neurons. While on average, T_in_ neurons exhibit a stereotyped level of activity before a contralateral choice, many neurons depart from this average in a consistent manner. From these neurons, the accuracy of the choice can be predicted using a simple logistic decoder. The accuracy of the choice predicted from neural activity reproduces the hallmarks of confidence identified in human behavioral experiments. Therefore, neurons that represent evidence accumulation can also inform the monkey’s confidence.

## Introduction

Choice, reaction time, and confidence are often considered the three pillars of choice behavior; a comprehensive model of decision-making should account for all three. However, the most prevalent models of binary decision making, Signal Detection Theory (SDT) and the Drift-Diffusion Model (DDM), can only account for two. In SDT, the choice depends on the sign of the difference between a sample of evidence and a decision criterion; confidence is a monotonically increasing function of the magnitude of that difference (Galvin et al., 2003; Kepecs and Mainen, 2012). Because SDT frames the decision as a categorization of one sample of evidence, SDT cannot account for reaction times, other than to posit that evidence closer to criterion might lead to slower choices, linked to uncertainty (e.g., Carpenter and Williams, 1995). In the DDM, the decision is made by accumulating many samples of evidence over time (Ratcliff, 1978; Palmer et al., 2005; Shadlen et al., 2006; Ratcliff et al., 2016). The decision ends when the accumulated evidence exceeds an upper or lower bound. The model naturally accounts for choice and reaction time, but lacks a straightforward explanation of confidence. This is because at the moment of choice, the state of accumulated evidence is uninformative about the accuracy of the choice.

In the brain, simple binary decisions are implemented as a race between two competing evidence-accumulation processes (Gold and Shadlen, 2007; Hanks et al., 2015). The first process to reach an upper bound terminates the decision and determines the choice and reaction time (RT). A case in point is the random dot motion task, in which monkeys make binary decisions about the net direction of random dot motion and communicate their decision with a saccadic eye movement. Neurons in the lateral intraparietal area (LIP) with response fields overlapping the choice-target contralateral to the recording site (T_in_ neurons) represent the accumulation of evidence in favor of contralateral target selection (Steinemann, Stine et al., 2024; Roitman and Shadlen, 2002). Related neural responses have been found in other brain areas and species (Hanks et al., 2015; Horwitz et al., 2004; Ding and Gold, 2010; Thura et al., 2012; Yartsev et al., 2018; Steinmetz et al., 2019). The decision process is well captured by race models of decision-making, which generalize drift-diffusion models by allowing the competing evidence-accumulation processes to be imperfectly anticorrelated (Wang, 2002; Usher and McClelland, 2001). On trials where the monkey chooses the contralateral target (i.e., the one within the neurons’ response field), the T_in_ neurons represent the *winning* race, whereas when the monkey chooses the ipsilateral target, the same neurons represent the *losing* race.

In addition to being supported by neurobiology, race models offer the leading explanation of choice, reaction time, and confidence in simple binary decisions, guided by the accumulation of evidence. Confidence is often modeled under the *balance of evidence* hypothesis, which postulates that confidence is a function of the difference, at the moment of choice, between the evidence accumulated by the winning race and the losing race (Vickers, 1979). Since the evidence accumulated by the winning race at the moment of choice is at its upper bound, confidence is determined by the state of the accumulation that has not reached its upper bound (i.e., the losing race). The greater the distance is from its upper bound, the weaker the accumulated evidence is for the losing alternative, and the stronger the confidence is for the chosen alternative. Models that embrace the balance of evidence hypothesis are able to account for several behavioral regularities of confidence, including the relationship between confidence and evidence strength, reaction time, and accuracy (Vickers 1979; Kiani et al. 2014; van Den Berg et al. 2016; Brus et al. 2021; Smith and Vickers 1988; Hellmann et al. 2023; Moreno-Bote 2010; Rolls et al. 2010; Wei and Wang 2015; Vivar Lazo 2024; Vickers et al. 1985, but see Zylberberg et al. 2012; Comay et al. 2023).

A less explored possibility is that confidence may depend not only on the relative difference between the state of scalar drift-diffusion processes but also on systematic variation within the neural populations that collectively manifest these processes. Each competing drift-diffusion process presumably represents the average activity over a large population of neurons with similar response properties (Steinemann, Stine et al., 2024). Theoretical studies have proposed that variability in firing rates within the population of neurons supporting the winning race may be used to inform confidence (Paz et al., 2016). Thus, rather than reflecting the difference between scalar decision variables, confidence may exploit within-population variability among neurons with shared choice preferences (Paz et al., 2016; Maniscalco et al., 2021; Fan et al., 2024).

We tested a key prediction of balance of evidence models, namely, that the state of the losing race at decision termination contains more information about the accuracy of the choice than the state of the winning race. We reanalyzed data from a recently published study. Steinemann, Stine et al. (2024) used high-density Neuropixels probes to obtain simultaneous recordings from large populations of neurons in macaque area LIP. Two monkeys made perceptual decisions about the net direction—left or right—in a stochastic random dot motion display and communicated their decision with a saccadic eye movement to one of two choice targets. We use the single-trial population response of T_in_ neurons to predict the accuracy of the choice, focusing on a brief (100 ms) window preceding the saccadic choice-report. Contrary to the prediction of balance of evidence models, information about the accuracy of the choice is more strongly encoded by neurons representing the winning race than by those representing the losing race. We show that these accuracy predictions also exhibit signatures of confidence reports observed in human behavioral studies.

These findings are unexpected because the average firing rate of T_in_ neurons is known to reach a stereotyped level just before the monkey issues a saccadic eye movement to the contralateral choice target. The key insight is that the average firing rate of the T_in_ neurons belies considerable heterogeneity of T_in_ responses across the population. We find that not all T_in_ neurons reach a common level of activity at the time of choice. Instead, some maintain a trace of evidence strength, while others maintain a representation of elapsed decision time. Through clustering analysis, we demonstrate that information relevant to encoding choice accuracy is distributed across two sub-populations of T_in_ neurons. These sub-populations carry information about the strength of the evidence supporting the choice and response time. Our findings suggest that the heterogeneous activity of T_in_ neurons during the final 100 ms of the decision process provides sufficient information for a linear readout of confidence that the choice is correct.

## Results

### Task, behavior and neurophysiological recordings

We analyzed previously published Neuropixels recordings from area LIP of two rhesus monkeys (*Macaca mulatta*), trained to make perceptual decisions about the net direction of motion in a dynamic random dot display (Steinemann, Stine et al., 2024). The monkeys indicated their choices by redirecting their gaze from a central fixation point to a left or right choice target. Monkeys were allowed to indicate their decision when ready, thus giving rise to two behavioral measures, choice and reaction time^1^ (RT) (Fig. 1A). The degree of difficulty was controlled by the *motion coherence*, defined as the probability that a dot displayed at time *t* will be redrawn in the direction of motion when replotted 40 ms later, as opposed to being randomly repositioned. On each trial, motion coherence was selected pseudorandomly from the list {±0%, ±3.2%, ±6.4%, ±12.8%, ±25.6%, ±51.2%}. The sign of the motion coherence indicates the direction (positive for leftward). For the 0% coherence motion, the sign indicates the random direction to be rewarded on that trial. The proportion of leftward choices increases with motion coherence, and RT shortens as a function of motion strength (the absolute value of motion coherence) (Fig. 1B). On about half of the trials, a brief (100 ms) pulse of motion, equivalent to a small change in motion coherence, was presented at a random time.

**Figure 1.**
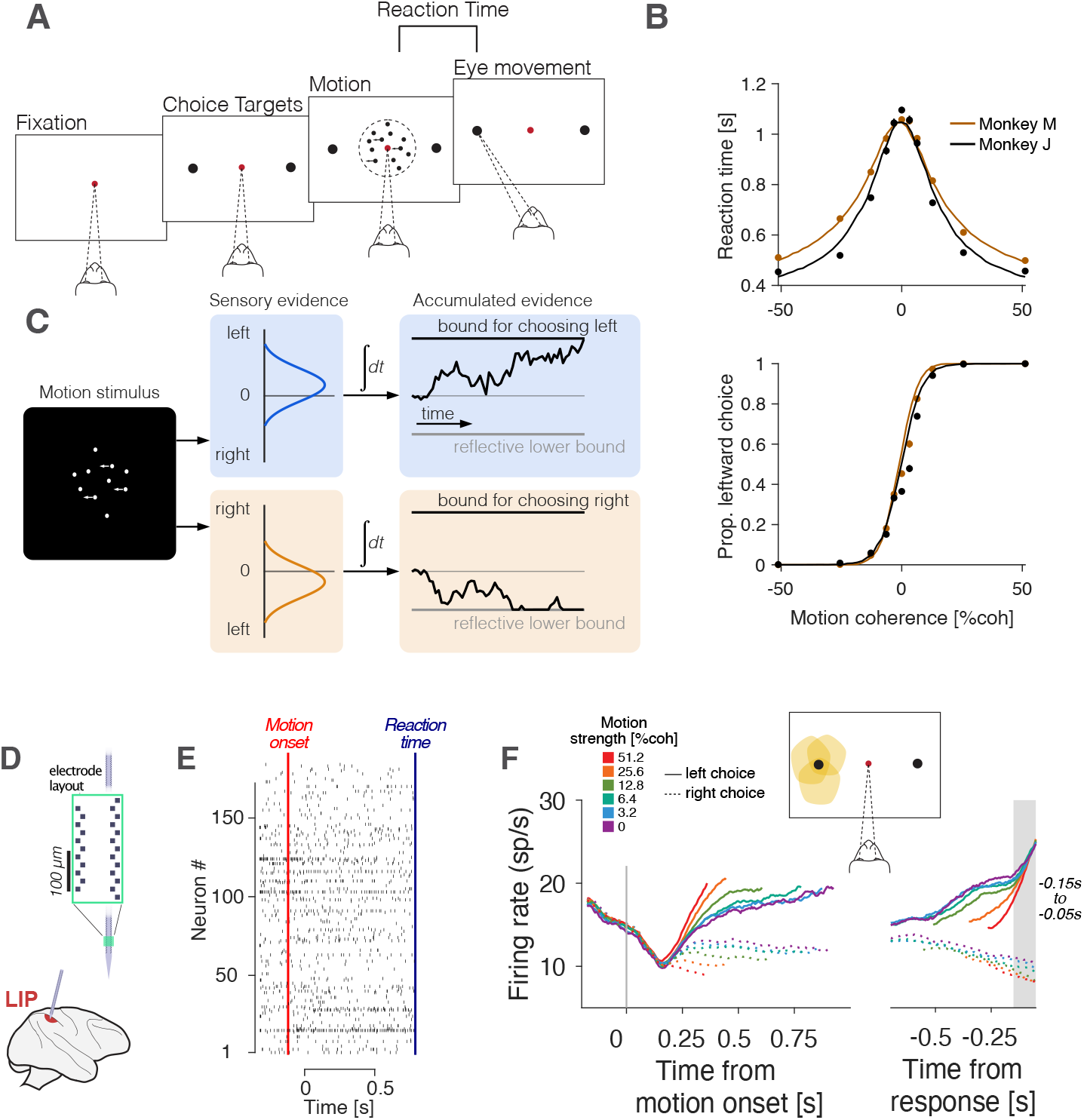
Large-scale recordings from LIP in a decision-making task. **(A)** Sequence of events in the random dot motion task. After the monkey fixates on a central spot, two choice targets are displayed, followed, after a random delay, by the random dot motion stimulus. The monkey is free to report its decision when ready by making a saccadic eye movement to one of the choice targets. The monkey is rewarded for choosing the left or right target for leftward or rightward motion, respectively. On noise-only (0% coherence motion) trials a reward is given with probability 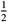. **(B)** Psychometric functions for the two monkeys studied by Steinemann, Stine et al. (2024). The average reactio^2^n time (top), and proportion of leftward choices (bottom) are plotted as a function of motion coherence. Positive and negative coherence indicate leftward and rightward motion, respectively. The solid lines are fits of the race model depicted in the next panel. **(C)** Sketch of the race model. The random dot motion stimulus provides sequential samples of momentary evidence for right minus left and left minus right, which are accumulated as a function of time to render two drift-diffusion processes. The samples are idealized as draws from Normal distributions with means proportional to motion coherence. The samples are anticorrelated (*ρ* = −0.7). The first process that reaches its positive bound terminates the decision and resolves the choice and decision time. A lower non-absorbing bound constrains the values of the negative accumulation. The reaction time is the sum of the decision time and a normally distributed non-decision time. **(D)** Schematic representation of the Neuropixels probe used to record neural activity from LIP in the right hemisphere (both monkeys). **(E)** Raster plot showing the spiking activity of 191 neurons simultaneously recorded during a representative trial. The red and blue vertical lines indicate the onset of motion and the choice report, respectively. **(F)** Average response of T_in_ neurons (N = 152) aligned with motion onset (left) and saccade initiation (right). Motion strength is indicated by color (legend). Solid lines indicate trials with leftward motion and dashed lines indicate trials with rightward motion. Only correct trials are included. The gray shading indicates the time between 150 ms and 50 ms before saccade initiation; most of our analyses focus on this time period.

The relationship between choice, reaction time, and motion coherence is well captured by a race model in which two drift-diffusion processes compete until one of them reaches a threshold or bound (Fig. 1C). The first process that reaches its upper bound determines the choice and the decision time. The reaction time is the sum of the decision time and a non-decision time, which is assumed to be Normally distributed and independent of decision time. In our instantiation of the race model, the drift-diffusion processes cannot fall below a lower *reflective* bound, which realizes the constraint that firing rates are non-negative (Zylberberg and Shadlen, 2016).

In addition to the main task, the monkeys also made visually-guided and memory-guided saccades to peripheral targets after variable delays (see Methods) (Gnadt and Andersen, 1988; Mazzoni et al., 1996; Colby et al., 1996). In the memory-guided saccade task, a target is briefly presented in the periphery while the monkey maintains its gaze on a fixation point. When the fixation point is extinguished, the monkey makes an eye movement to the remembered location of the target. The task served to identify, *post hoc*, neurons that display persistent activity as the monkey plans a saccadic eye movement towards the choice target contralateral to the recording site. These neurons are referred to as T_in_ neurons. Monkeys also performed a passive motion viewing task in which they were rewarded for maintaining fixation while viewing random dot motion (higher motion strengths only; see Methods).

The neural data from Steinemann, Stine et al. (2024) and Stine et al. (2023) were recorded from area LIP using high-density NHP-neuropixels probes. Between 54 and 203 single neurons were recorded simultaneously over eight recording sessions (mean = 135.5 neurons/session) (Fig. 1E). Steinemann, Stine et al. (2024) showed that T_in_ neurons represent the drift-diffusion signal associated with stochastic choice and reaction time on single trials. On average, T_in_ neurons tend to ramp with positive slope on trials where the monkey chooses the target in the neurons’ response field, and this slope is steeper as a function of motion strength (Fig. 1F)(Roitman and Shadlen, 2002). Importantly, the population of T_in_ neurons reach a common level of activity ∼ 100 ms before a saccadic eye movement towards the contralateral choice target. These observations conform to the predictions of the race model illustrated in Fig. 1C, under the assumption that the T_in_ neurons represent the accumulation of evidence for a contralateral choice, and that another (unobserved) population of neurons represents the accumulation of evidence for the ipsilateral (rightward) choice (e.g., Usher and McClelland, 2001; Wong and Wang, 2006).

### Choice accuracy decoded from LIP population activity

We investigated whether T_in_ neurons, which represent the accumulation of noisy evidence, or drift-diffusion, are also predictive of whether the choice would be the correct one and thus capable of informing the monkey’s confidence (or reward prediction). We train a logistic decoder to predict the accuracy of each contraversive (i.e., left) choice from the activity of the T_in_ neurons shortly before the choice is reported:

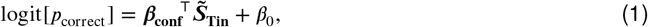

where ***β***_**conf**_ is a column vector of regression coefficients with as many elements as there are T_in_ neurons in the session, and *β*_0_ is a bias term.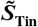 contains the standardized (i.e., z-scored) number of spikes emitted by each T_in_ neuron in the time interval between 150 ms and 50 ms before the choice report; we refer to this time interval as the *presaccadic window* throughout the manuscript. The model is fit separately for each session and choice category, left or right, including both correct and error trials. The logistic decoder yields a probability that quantifies the confidence the decision-maker should have in the choice. The model is fit separately for each session using 10-fold cross-validation. Specifically, we divide the data into 10 groups of an approximately equal number of trials, using one group as a prediction set, and the remaining 9 for training; we repeat this process 10 times so that confidence estimates for every trial are based on a prediction.

We use a receiver operating characteristic (ROC) analysis to assess how effectively the probability correct, predicted with Eq. 1, distinguishes between correct and incorrect choices. The approach is illustrated in Fig. 2A. The figure shows the predicted probability correct for trials with a contraversive (i.e., left) choice. Correct and incorrect decisions are indicated in blue and red, respectively. The area under the ROC curve (AUC_conf_) is a measure of how well the predicted probability correct for each trial discriminates correct from incorrect choices (Fig. 2A, inset). More specifically, it is the probability that given two choices, one correct and one incorrect, the predicted probability correct is greater for the correct choice. Using this simple metric we can evaluate the balance of evidence hypothesis mentioned above.

**Figure 2.**
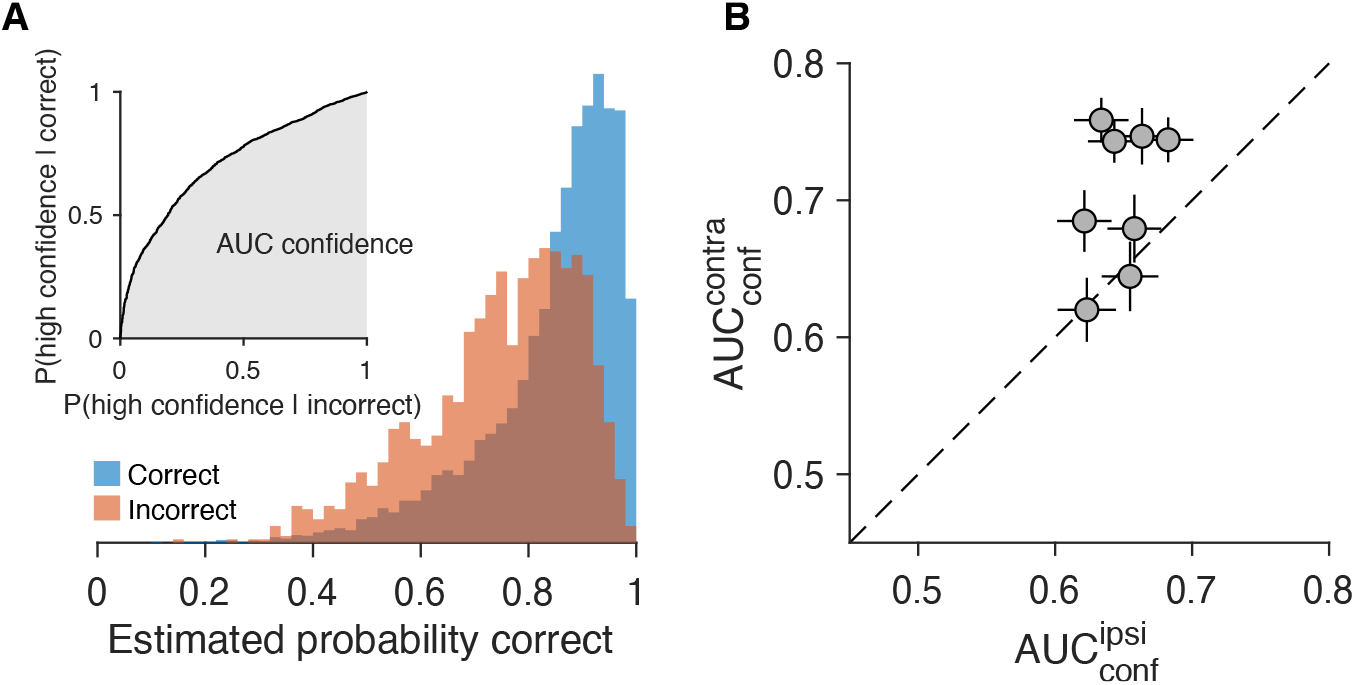
Confidence inferred from the population of T_in_ neurons. **(A)** Confidence estimates obtained with Eq. 1, shown separately for factually correct (blue) and incorrect (red) choices. These distributions are used to construct an ROC (inset) and to calculate the area under the ROC (AUC_conf_; gray). Confidence estimates from the eight sessions are pooled. Only trials with contralateral (left) choices are included. **(B)** Area under the confidence ROC calculated from trials in which the ipsilateral (abscissa) or contralateral (ordinate) target was chosen. Each data point corresponds to a different session. Error bars indicate s.e. (bootstrap).

We compared the AUC_conf_ derived from the logistic model fit separately for contralateral and ipsilateral choices. As mentioned, T_in_ neurons represent the winning race for contralateral choices and the losing race for ipsilateral choices. Therefore, according to the balance of evidence hypothesis, the T_in_ neurons should contain more information about choice accuracy when monkeys select the ipsilateral (right) target, than when they select the contralateral (left) target. Fig. 2B shows the AUC_conf_ for each of the 8 recording sessions. Contrary to the balance of evidence hypothesis, T_in_ neurons contain more information about choice accuracy when the monkey chooses the contralateral target (i.e., when the T_in_ neurons represent the winning race) than when it chooses the ipislateral target (*p* = 0.008, one-tailed *t-test*).

As mentioned earlier, this finding is surprising because the T_in_ neurons appear to reach a stereotyped level of activity before a contralateral choice, independent of motion strength and reaction time (Fig. 1F; Roitman and Shadlen 2002). Since motion strength and reaction time are strong predictors of accuracy, one would not expect T_in_ neurons to contain information about choice accuracy in the presaccadic window. Indeed, this is the central premise of the balance of evidence hypothesis. We will address this tension after bolstering the claim that the choice accuracy that we infer from neural activity replicates the behavioral hallmarks of confidence identified in human psychophysical experiments.

### Choice accuracy inferred from neural activity reproduces behavioral features of confidence

The monkeys did not report their confidence, so we cannot establish a correlation between the putative confidence signal and behavior. Instead, we ask whether a monkey exploiting this signal would mimic the regularities of confidence reports observed in humans performing a task similar to the one performed by the monkeys. van Den Berg et al. (2016) asked human participants to perform a variant of the random dot motion task in which they reported choice and confidence (high/low) simultaneously by moving a handle to one of four targets (Fig. 3A). Results from a representative participant are shown in Fig. 3A. The data show that: (*i*) confidence is greater for correct than for incorrect decisions, even when controlling for motion strength (Fig. 3A, left), (*ii*) for correct decisions, confidence increases as a function of motion strength (Fig. 3A, left), (*iii*) for incorrect decisions, confidence also increases as a function of motion strength (Fig. 3A, left), (*iv*) confidence decreases as a function of reaction time (for both correct and incorrect decisions) (Fig. 3A, center), (*v*) for a given reaction time, confidence is lower for incorrect decisions than for correct decisions (Fig. 3A, center), (*vi*) for a given reaction time, confidence increases as a function of motion strength, even when controlling for accuracy (Fig. 3A, right).

**Figure 3.**
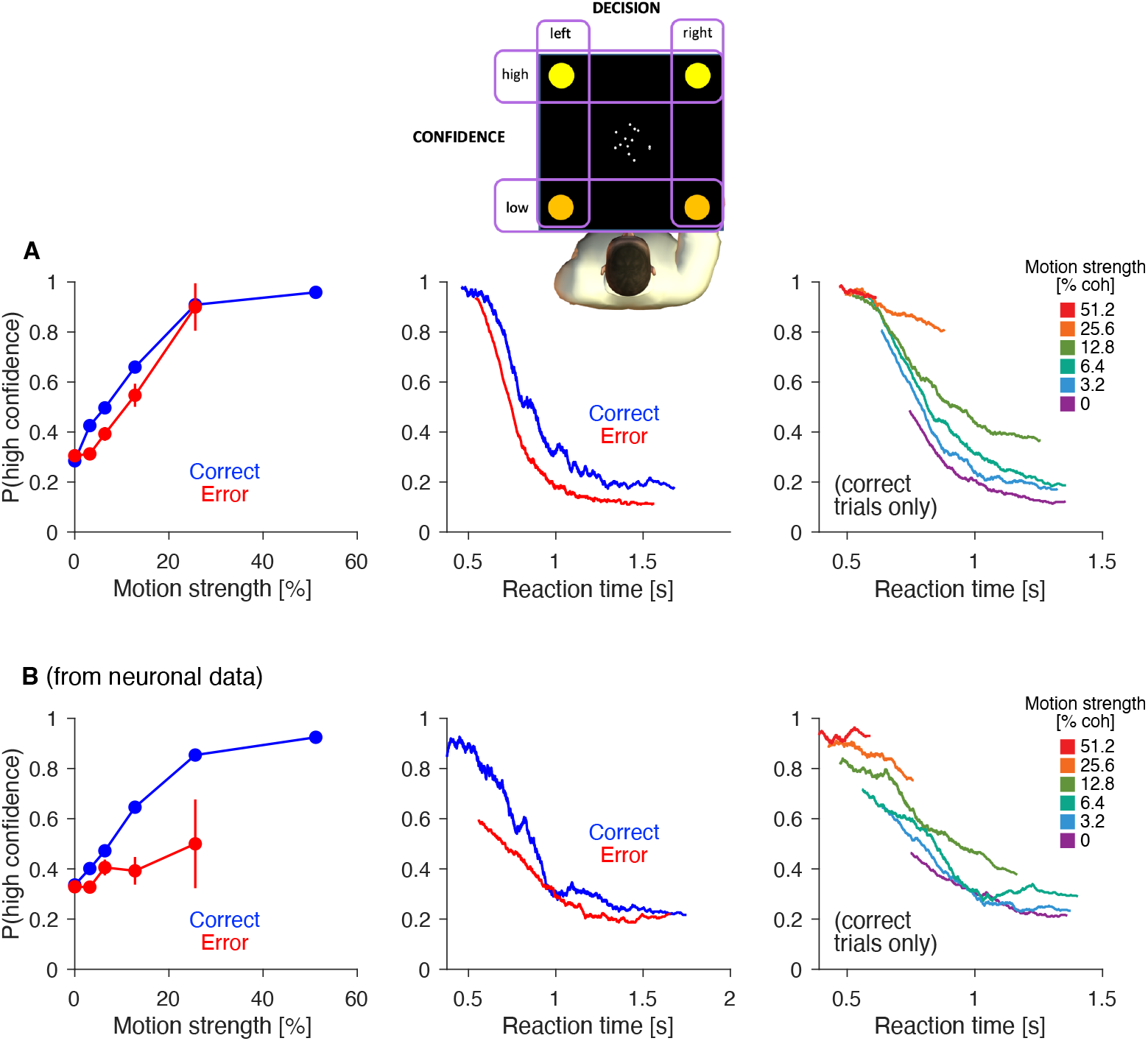
Choice accuracy inferred from neural activity reproduces behavioral signatures of confidence. **(A)** Random dot motion task with simultaneous choice and confidence reports (from van Den Berg et al., 2016). The two left and right targets are used to indicate leftward and rightward motion. In alternating blocks, either the top two targets or the bottom two targets were used to indicate high-confidence choices, and the remaining two targets were used to indicate low-confidence choices. Data correspond to a representative participant from van Den Berg et al. (2016) (N = 9,024 trials; proportion of high-confidence choices: 61%). Data from the other participants in van Den Berg et al. (2016) are shown in Fig. S1. *left*. Proportion of high confidence choices as a function of motion strength, shown separately for correct and incorrect choices. Conditions with fewer than 4 trials were excluded. *center*. Proportion of high confidence choices as a function of reaction time, shown separately for correct and incorrect choices. Trials were sorted by reaction time and smoothed with a boxcar filter (N = 300 trials). *right*. Proportion of high confidence choices as a function of reaction time, for correct trials only, plotted separately for each motion strength. Trials were sorted by reaction time and smoothed with a boxcar filter (N = 300 trials). **(B)** Same analyses as in panel A, but for the Steinemann, Stine et al. (2024) data, using as confidence the probability of correct inferred from the population of T_in_ neurons using logistic regression (Eq. 1). The continuous confidence estimate was thresholded so that the proportion of trials with high confidence was the same as in the behavioral data (panel A). A version with non-thresholded confidence estimates is included as Fig. S2A.

The putative confidence signal reproduces these observations. To parallel the design of van Den Berg et al., we thresholded the putative confidence signal (obtained with Eq. 1) using a criterion set such that the proportion of high-confidence choices is equal to the proportion of high-confidence reports in the human experiment (61% high-confidence choices). Without any free parameters, the confidence signal qualitatively reproduces all the behavioral hallmarks of confidence observed by van Den Berg et al. (Fig. 3B). The similarity is not a consequence of the thresholding step in the analysis. The same qualitatve reproduction of the human regularities are also present without thresholding (Fig. S2).

In short, the population activity of T_in_ neurons measured just before a contralateral choice report contains information bearing on whether the choice is correct or incorrect. The confidence signal varies with motion strength, reaction time, and accuracy, in a similar manner to explicit confidence reports. Importantly, it is not necessary to consider the state of the losing race or the decision time to reproduce the behavioral features of confidence; the necessary information is contained in the population response of the T_in_ neurons.

Because of the tight link between confidence and reaction time (e.g., Henmon, 1911), we reasoned that the population activity of T_in_ neurons just before the response should contain information about RT. We fit a regression model to the reaction times for trials with a contralateral choice, again using the spike counts of the T_in_ neurons in the presaccadic window (Eq. 8). If all T_in_ neurons reach a stereotyped level of activity before the response, then there would be no information about reaction time just before the choice. Contrary to this prediction, we were able to distinguish longer from shorter reaction time (relative to the median) reliably from the population activity of T_in_ neurons (AUC= 0.85 ± 0.02; mean ± s.e. across sessions). Thus, shortly before the choice report T_in_ neurons contain information about the time required to make the decision. This information likely contributes to the ability of the accuracy decoder to reproduce the behavioral features of confidence that are thought to require an explicit representation of decision time.

We conducted a control analysis to assess whether the prediction of choice accuracy from neural activity could be attributed to differences in the amplitude of the eye movement used to report the choice. This scenario seemed unlikely a priori, as the epoch ends 50 ms before saccade initiation, when the monkey is still holding fixation. To test this, we repeated the decoding analysis, substituting the neural signals in Eq. 1 with the X and Y coordinates of the gaze averaged over two intervals: 100 ms before saccade detection and 100 ms after saccade completion (yielding four regressors). The capacity to decode choice accuracy from these signals was significantly lower than that achieved with T_in_ neurons (AUC_conf_ = 0.57 ± 0.01 vs. 0.72 ± 0.01, respectively; *p*<10^−6^, bootstrap). Furthermore, this predicted accuracy from the gaze signals did not reproduce the behavioral characteristics of confidence (Fig. S2B).

### Heterogeneity of T_in_ responses underpins the representation of choice accuracy

The observation that these neurons achieve a stereotyped state at the end of a contralateral choice would appear to be incompatible with a graded representation of choice accuracy. However, this stereotyped state is evident in firing rates averaged over many neurons (e.g., Roitman and Shadlen, 2002; Steinemann, Stine et al., 2024). We thus considered the possibility that the population of T_in_ neurons might contain information about choice accuracy that is lost in the firing rate averages.

To test this possibility, we used a combination of linear regression and *k-means* clustering. The linear regression sets out to explain the spike counts in the presaccadic window for each T_in_ neuron on each trial, using three variables: the motion coherence, choice, and reaction time, plus an offset (Eq. 9). We fit the model independently for each neuron and applied *k-means* to assign the neurons to three clusters. This classification was based on the regression coefficients associated with the three variables (Fig. 4A).

**Figure 4.**
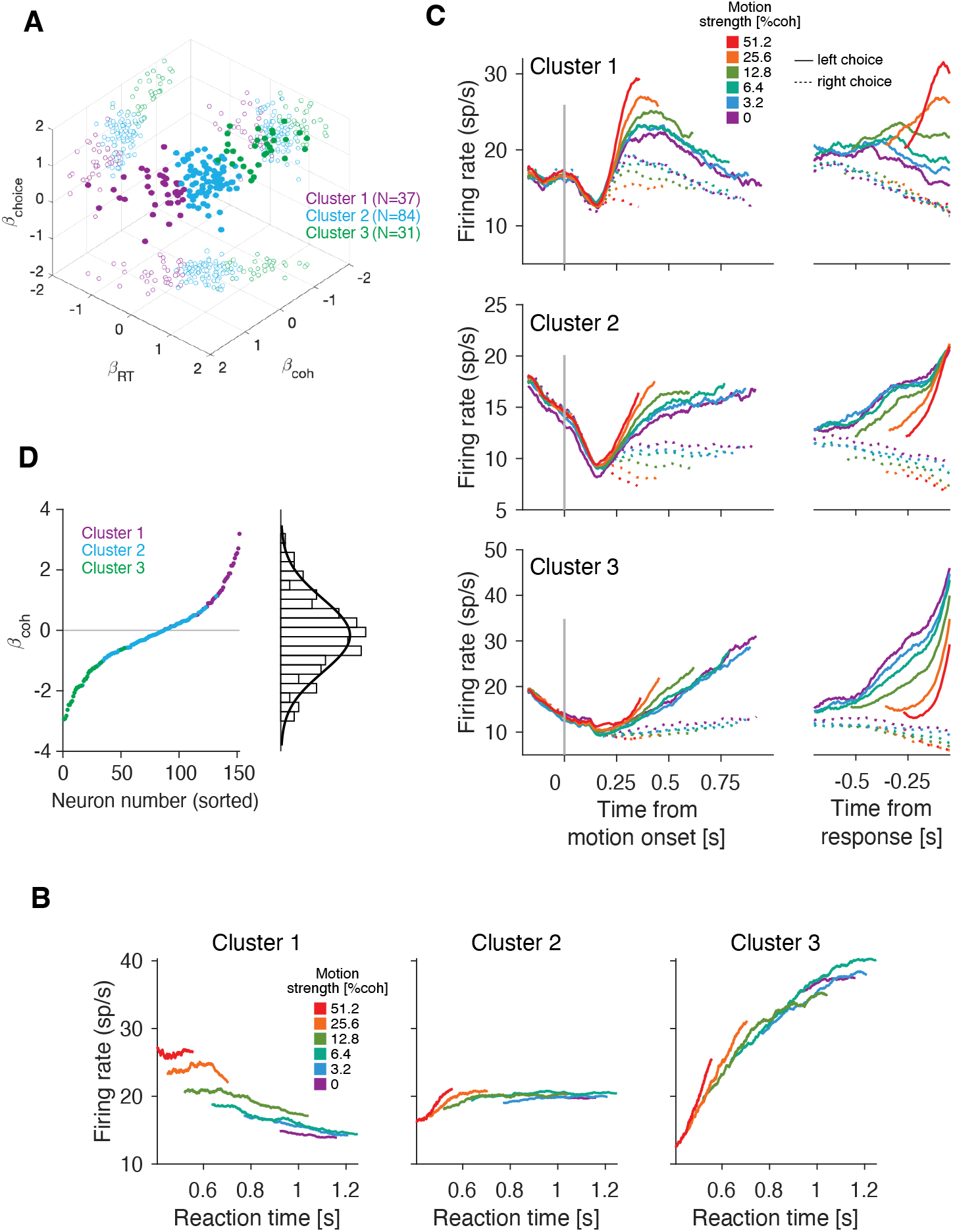
Distinct response characteristics of T_in_ neurons. The T_in_ neurons vary in their representation of motion coherence, reaction time and choice. We distinguished three clusters, based on a linear regression model of these explanatory variables on each T_in_ neuron’s response. **(A)** Regression coefficients associated with choice, motion coherence, and reaction time for each neuron. Each filled symbol shows the 3-tuple of one neuron. Color shows the cluster membership assigned by *K-means* procedure instructed to identify three clusters. Open circles are the 2D projections of the regression coefficients. The number of T_in_ neurons in each cluster is shown in parenthesis. **(B)** The average firing rate of the neurons within each cluster, calculated within the presaccadic window, is plotted as a function of reaction time. Traces are calculated separately for each motion strength, including only correct contralateral choices. Traces are smoothed with a boxcar filter (N = 500 trials). **(C)** Average response of T_in_ neurons for each cluster. Same conventions as in Figure 1F. **(D)** Regression coefficients associated with motion coherence, obtained from a linear regression using motion coherence (plus an offset) to explain the spike counts (z-scores) emitted by each neuron in the presaccadic window. The colors indicate the cluster to which each neuron belongs. The histogram of regression weights (right) is well described by a Normal distribution (thick black trace) with a mean and standard deviation of −0.17 and 1.2, respectively.

The T_in_ neurons cluster into groups that exhibit distinct response characteristics. Figure 4B shows the firing rates of neurons within each cluster, calculated within the presaccadic window, as a function of reaction time and split by motion strength. The analysis includes only correct contralateral choices. Just before the choice is reported, neurons in cluster 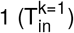 exhibit higher firing rates for strong motion and faster choices. The traces corresponding to different motion strengths do not converge when conditioned on reaction time, indicating that the activity of these neurons at the moment of choice is informative about both reaction time and motion strength. In contrast, neurons in cluster 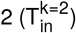 appear to be largely unaffected by reaction time or motion strength, consistent with expectations for neurons that reach a stereotyped level of activity prior to response. For the neurons in cluster 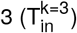 the activity increases strongly with reaction time and is only slightly influenced by motion strength.

We repeated the clustering analysis using only the odd-numbered trials or only the even-numbered trials from each session. The regression weights assigned to motion coherence, RT and choice were highly consistent across both analyses (Fig. S3). Applying *k-means* to the coefficients derived from odd and even trials, we observed that most neurons (89%) were assigned to the same cluster in both realizations, indicating that the neuron clustering is robust.

Fig. 4C compares and contrasts the response properties of the three clusters. The graphs show the mean firing rate of neurons within each cluster, aligned to motion onset and reaction time. Neurons from all three clusters are choice selective; compare solid and dashed traces in Fig. 4C and the responses on correct and error choices (Fig. S5A). Neither 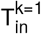 nor 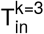 neurons reach a common level of activity prior to the response.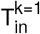 neurons do not show the ramping activity usually associated with T_in_ neurons. Instead, the traces for the different motion coherences are largely parallel to each other, similar to the representation of momentary evidence in upstream visual areas (e.g., Britten et al., 1992). For contralateral choices, 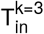 neurons appear to increase their firing rate more rapidly than 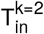 or 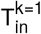 neurons. The firing rate of 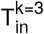 is higher just before choice in difficult (i.e., low motion strength) decisions compared to easier ones.

The latency to choice selectivity was similar across clusters. We calculated this latency independently for each neuron using the CUSUM method (Ellaway, 1978; Lorteije, Zylberberg et al., 2015). The average latency to direction selectivity was 0.22±0.03, 0.22±0.02 and 0.23±0.03 s from motion onset for neurons in clusters 1, 2, & 3, respectively. No significant differences in latency were observed between clusters (Fig. S4) (*p*>0.3 for all three pairwise comparisons, *t-test*).

The heterogeneity of neuronal responses across the population of T_in_ neurons can also be observed by analyzing individual neurons, without clustering. We used a linear regression model to characterize the relationship between motion coherence and the neuronal activity in the presaccadic window. In the regression model, motion coherence for each trial (plus an offset) was used to explain the standardized (z-scored) spike counts in the presaccadic window. Only trials with a contralateral choice were included. The regression coefficient associated with motion coherence varies substantially between neurons (Fig. 4D). For some neurons, activity increases with motion coherence (*β*_coh_>0), while for others, it decreases (*β*_coh_<0). The values of *β*_coh_ are approximately normally distributed (Fig. 4D). Because the mean is close to zero, the activity of T_in_ neurons just before the response appears to be unaffected by motion coherence when averaged over many neurons (Fig. 1F; Roitman and Shadlen 2002).

### The heterogeneity of neuronal responses is not evident in control tasks

The memory-guided saccade task was used to identify the T_in_ neurons and historically to elucidate the their hallmark visual, memory and perisaccadic responses (Gnadt and Andersen, 1988; Mazzoni et al., 1996; Colby et al., 1996). We wondered whether the signs of the heterogeneity we identified in the random dot motion task would be evident in the memory-guided saccade task. Fig. S5B shows the response of neurons from the three clusters for memory saccades to the contralateral and ipsilateral targets. The three clusters show similar persistent activity during the delay period. We calculated the spike counts in the last 200 ms before the fixation point is extinguished, and computed the average difference in standardized counts between trials with memory-saccades to the left and right target. This measure fails to differentiate the three clusters (*p*_max_>0.2, Wilcoxon rank–sum test).

The monkeys also performed a passive motion viewing task in which they were presented with strong (*c* = ± 51.2%) left and right motion and were rewarded for maintaining fixation. We expected neurons in cluster 1 that retain information about motion strength to have response fields that overlap the random dot motion stimulus, but there is no sign of this. Neurons in all three clusters maintained low activity during the passive viewing task (Fig. S5C). While firing rates were significantly greater for leftward than for rightward motion (p = 0.047, 0.0026, and 0.0145 for clusters 1, 2, and 3, respectively; one-sided Wilcoxon signed-rank test), decoding accuracy was poor and similar so for all clusters. We calculated the spike rate during motion viewing (excluding the initial 200 ms), and computed the average difference in standardized rates between trials with leftward and rightward motion. This measure does not distinguish between the three clusters (*p*_max_>0.25, Wilcoxon rank–sum test). The capacity of this measure to distinguish between leftward and rightward motion was low (AUC =0.55 ± 0.01, mean ± s.e. across neurons). As an additional control, we repeated the decoding and clustering analyses after excluding the T_in_ neurons that significantly discriminated between leftward and rightward motion in the passive motion–viewing task. Excluding these neurons (N = 45) yielded qualitatively similar results Fig. S6. We conclude that the response features that distinguish the three clusters of T_in_ neurons are not elucidated by memory saccades or passive motion viewing. Instead, the discriminating feature is more likely associated with the decision mechanism.

### Clusters 1 and 3 are the most informative about the accuracy of the choice

Because confidence is informed by evidence strength and decision time, and because these variables are represented by the population of cluster 1 and cluster 3 neurons, we reasoned that neurons from these clusters should be the most informative about choice accuracy. We repeated the logistic regression analysis used to decode accuracy from population activity (Eq. 1), using signals from one cluster at a time. Neurons in cluster 1 and cluster 3 are more predictive of choice accuracy than those in cluster 2 (Fig. 5A) (*p*_max_<10^−8^, bootstrap).

**Figure 5.**
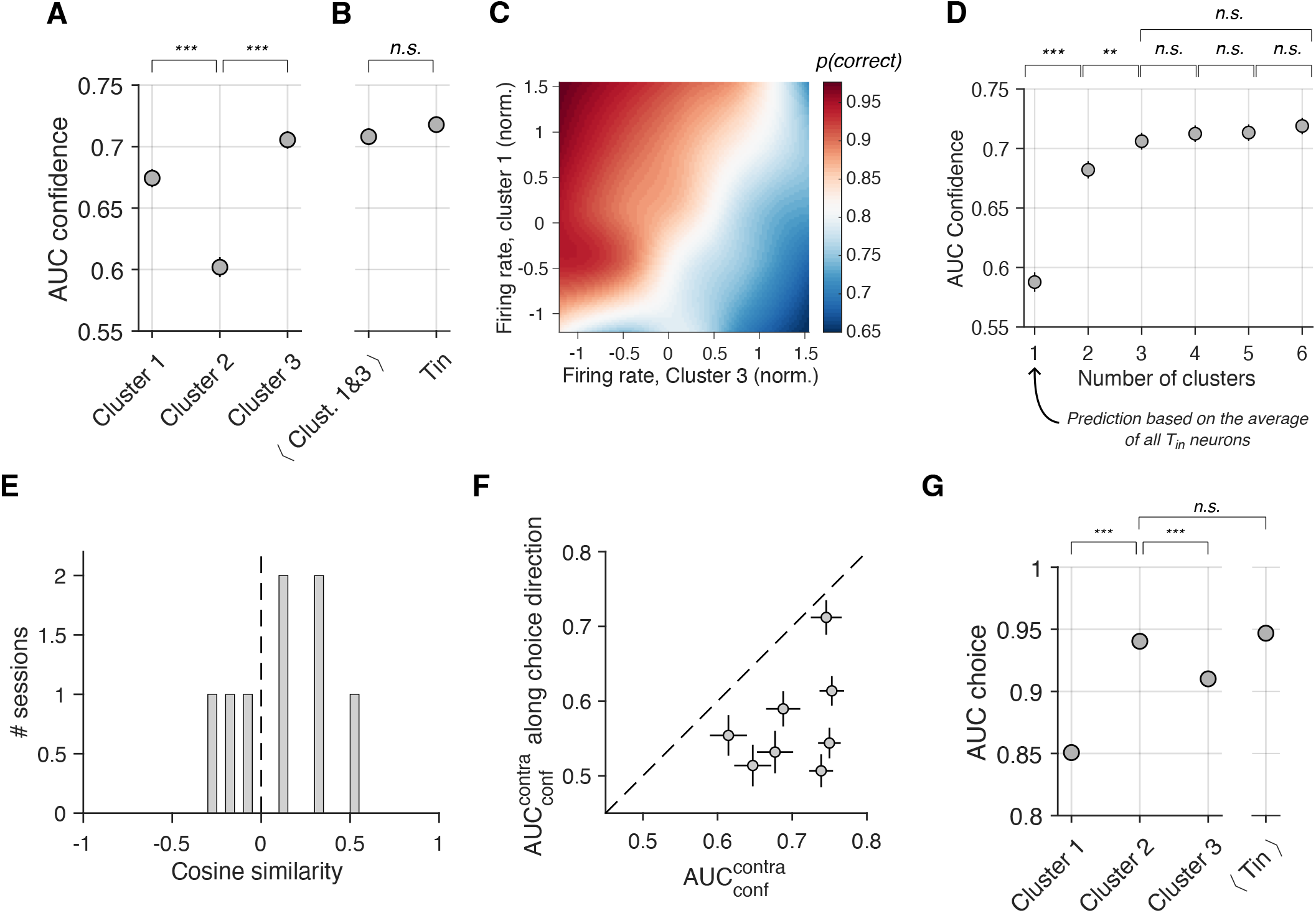
Information about choice accuracy differs across clusters. **(A)** Neurons from cluster 3 are the most informative about choice accuracy. Symbols show the area under the receiver-operator curve (AUC_conf_) which quantifies the separation between the distribution of spike counts in the presaccadic window on correct vs. incorrect left choices. The accuracy predictions used to calculate the AUC_conf_ are based on the activity of the T_in_ neurons assigned to each of the three clusters (abscissa). Error bars indicate s.e. (bootstrap). Bootstrapping was used to assess statistical significance (n.s.: *p*>0.05; ***: *p*<10^−6^;**: *p*<10^−3^). **(B)** AUC_conf_ values obtained from the accuracy decoders trained on either (*i*) the average activity of neurons from cluster 1 and cluster 3, or (*ii*) the individual T_in_ neurons. **(C)** Proportion of correct responses calculated from the data as a function of the firing rate of neurons from cluster 1 (abcissa) and cluster 3 (ordinate). Spike counts in the presaccadic window from cluster 1 and cluster 3 neurons were independently z-scored within sessions, then grouped across sessions and divided into 5 percentiles. The proportion of correct responses was calculated for all combinations of percentiles and smoothed via cubic interpolation for finer resolution.**(D)**Clustering analysis applied to the regression coefficients obtained from Eq. 9, using different numbers of clusters (abccissa). We average the spike counts in the presaccadic window of neurons belonging to the same cluster. From these averaged spike counts, we calculate the AUC_conf_ (ordinate) using Eq. 6. We do not find a significant difference in the AUC_conf_ values between *N*=3 clusters and *N*>3 clusters (bootstrap). **(E)** Cosine similarities between the weights assigned by the regression to choice, ***β***_**choice**_, and the regression to accuracy, ***β***_**conf**_, across sessions. The set of weights define coding directions in the neuronal state space. **(F)** We project the neural activity onto the coding directions defined by ***β***_**choice**_ and by ***β***_**conf**_, and compute the information about choice accuracy, measured by the AUC_conf_, contained in these two projections. Information about choice accuracy is larger for the projection onto the direction defined by ***β***_**conf**_ (abscissa) than onto the one defined by ***β***_**choice**_ (ordinate). Each data point corresponds to a different session. Error bars indicate s.e. (bootstrap). **(G)** AUC_choice_ values derived from four separate regression analyses, incorporating neurons from clusters 1, 2, or 3 (left), or the average activity across all T_in_ neurons (right). Error bars represent standard errors (bootstrap). Neurons in cluster 2 are more informative about choice than those in clusters 1 or 3 but are equally informative as the average activity of all T_in_ neurons.

The average spike counts across the population of cluster 1 neurons, combined with the average across the population of cluster 3 neurons, predict choice accuracy with fidelity. We trained the accuracy decoder (Eq. 1) using two independent variables: the average of the spike counts from cluster 1 neurons and cluster 3 neurons within the presaccadic window (plus an offset). The AUC_conf_ from this regression model is statistically indistinguishable from the model using all T_in_ neurons (p = 0.16, bootstrap; Fig. 5B). By reducing the relevant information to two variables—the average spike counts of cluster 1 and cluster 3 neurons—we can visualize their relationship with the proportion of correct responses (Fig. 5C). The visualization reveals that higher cluster 3 activity and lower cluster 1 activity are associated with a greater proportion of correct choices. We conclude that the activity of cluster 1 and cluster 3 neurons (i.e., two numbers per trial) accounts for most of the information about choice accuracy contained in all recorded T_in_ neurons.

We chose to use three clusters for analytical convenience. The number is not guided by a biological or computational principle. Indeed, the regression coefficients used for clustering exhibit continuous variation (Fig. 4A). Nonetheless, three seems to be the right number, as increasing the number of clusters did not improve the prediction of accuracy, while decreasing the number of clusters diminished it. We repeated the decoding analysis using between one and six clusters. For each number, we calculated the AUC_conf_. We averaged the number of spikes in the presaccadic window across neurons belonging to the same cluster and used these averaged spike counts to predict choice accuracy (Eq. 6). We found a significant difference in AUC_conf_ between one and two clusters, and between two and three clusters, but not between three and four clusters or three and six clusters (Fig. 5D). Three clusters may be adequate for our purposes as they are the minimum number required to capture the central tendency and both signs of diversity. We emphasize, however, that we do not interpret this as evidence of three distinct neuron classes within the T_in_ population.

The average activity of T_in_ neurons reflects motion strength during the first half of the presaccadic window (Fig. 1F), which might raise concerns that the ability to distinguish correct from incorrect decisions based on the population response is primarily driven by this average activity. Three observations rule out this possibility: (i) when using a single cluster (*N*=1) for training the accuracy decoder, effectively relying on the within-trial average activity of all T_in_ neurons, the decoding capacity is low (Fig. 5D); (ii) excluding cluster 2 neurons—whose average activity closely resembles that of the full population—has little impact on the ability to decode choice accuracy (Fig. 5B); (iii) training the decoder using spike counts from the second half of the presaccadic window produces the same confidence signatures (Fig. S2C). These findings collectively demonstrate that the decoding capacity arises from more than just the average neuronal activity.

### Distinct contribution of T_in_ neurons to decoding choice and accuracy

We assessed whether the decoding of choice accuracy and the decoding of the choice itself relies on a common weighting of the activity of the T_in_ neurons. To this end, we fit a regression model similar to the one we used to predict decision accuracy (Eq. 1), but here the variable to predict is choice (left/right) (Eq. 7). We used the spike counts of the T_in_ neurons in the presaccadic window to derive the best-fitting regression weights, ***β***_**choice**_. The regression weights define a coding direction (CD) in the state space, where each T_in_ neuron represents a different dimension. Unsurprisingly, choice can be decoded with high precision from the population of T_in_ neurons (AUC_choice_ = 0.96 ± 0.014; mean ± s.e. across sessions). More interestingly, the cosine similarity between the directions defined by the regression weights on choice, ***β***_**choice**_, and the regression weights on accuracy, ***β***_**conf**_, is low: the average (across sessions) absolute value of the cosine similarity is 0.25 ± 0.05, indicating that the two directions in state space are closer to orthogonal than similar (Fig. 5E). Indeed, the projection of the population activity on the choice-CD is barely informative about accuracy (AUC_conf_ = 0.57 ± 0.024; mean ± s.e. across sessions) and significantly less informative than the projection on the confidence CD (p = 0.0009, *t-test*; Fig. 5F).

We reasoned that the distinct contributions of T_in_ neurons to decoding choice and accuracy may be evident at the level of clusters. We repeated the logistic regression analysis, again using signals from one cluster at a time (as in Fig. 5A) but now training the decoder to predict choice. As shown in Fig. 5G, the neurons from cluster 2 are more predictive of the choice than neurons from clusters 1 and 3 (*p*_max_<10^−8^, bootstrap). However, unlike what was observed with the accuracy decoder, the within-trial average activity of all T_in_ neurons (one count per trial) is just as informative as the decoding capacity of all neurons from the most informative cluster (Fig. 5G).

### Multiple time-scales of evidence accumulation represented by T_in_ neurons

We considered the possibility that the different response characteristics of T_in_ neurons might be explained by differences in how long (or persistently) the momentary motion evidence affects neuronal activity. We evaluate this idea by analyzing the influence of the short (100 ms) motion pulses introduced into the random dot motion stimulus. For each T_in_ neuron *i* and trial *j*, we count the spikes emitted between *t* and (*t*+100) ms, where *t* is the time from the onset of the motion pulse, and standardize the counts separately for each motion coherence and 100 ms time window. We refer to the standardized counts as 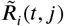. We then calculate the average difference, 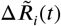, between trials with a leftward pulse and trials with a rightward pulse.

The difference 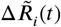 is a measure of the influence of the motion pulse on neuronal activity. We averaged this difference across neurons belonging to the same cluster and fit a function (Eq. 13) to these averages (Fig. 6A). The function implements two assumptions: (*i*) the pulse affects the neural response with a variable latency, and (*ii*) its effect dissipates exponentially with decay rate *α* (Eq. 13; Lorteije, Zylberberg et al. 2015). The best-fitting *α* values are 100, 3.1, & −0.17 for neurons of clusters 1, 2, and 3 respectively, consistent with a more persistent effect of the motion pulse on cluster 2 neurons than on cluster 1 neurons, and a more persistent effect on cluster 3 neurons than on cluster 2 neurons (Fig. 6A). The negative *α* (cluster 3) reflects the monotonic increase in the impact of the pulse as a function of time (Fig. 6A, *bottom*). We used a bootstrap analysis to create a distribution of *α* values for neurons of the three clusters. The probability of observing in the bootstrap distributions an *α* value greater in cluster 1 than in cluster 2 was 86.8 %, and a value greater in cluster 3 than in cluster 2 was 96.6 %. While the probability of observing this rank ordering by chance is not negligible, the analysis is consistent with the hypothesis that the time constant of evidence accumulation is different for neurons from the three clusters.

**Figure 6.**
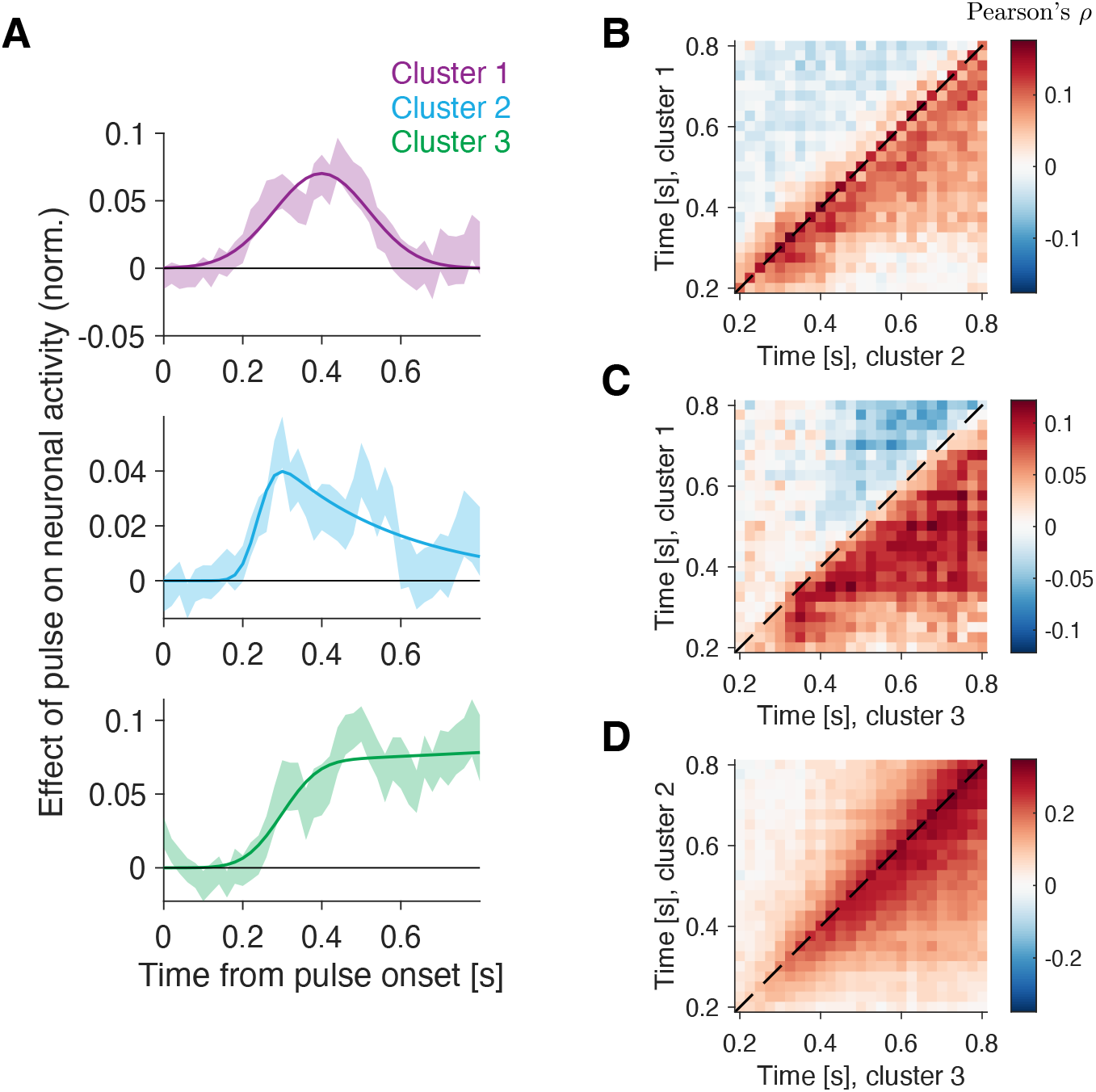
Different time constants of evidence integration by T_in_ neurons. **(A)**Influence of a brief motion pulse on the activity of cluster 1 (top), cluster 2 (middle) and cluster 3 (bottom) neurons (Eqs. 10 and 11). Error bars indicate s.e. (bootstrap). Solid lines are fits of a function (see Methods). The pulses affect the neural response with latency ≈ 200 ms. **(B–D)** Noise correlations between neurons from clusters 1 and 2 (panel B), clusters 1 and 3 (panel C), and clusters 2 and 3 (panel D). Time is relative to pulse onset. The correlation coefficients were calculated in non-overlapping 25 ms windows, using only trials with contralateral choices.

We further substantiate this interpretation with an analysis of the pairwise correlations between neurons belonging to different clusters. We counted the spikes of neurons from cluster 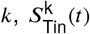, in 25 ms windows. We formed pairs { *x, y*}, where 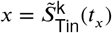 and 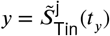, with *k* and *j* representing different clusters. The tilde in these expressions indicates the use of standardized residual values, for each motion coherence. The heat maps in Fig. 6B-D illustrate the correlation of these residuals across trials, for different pairs of clusters. Fluctuations in the activity of cluster 1 neurons predict—at later times— fluctuations in the activity of cluster 2 neurons, and the activity of cluster 3 neurons (both *p* < 10^−8^, permutation test; Fig. 6B & C). Similarly, a noise correlation analysis between the activity residuals of cluster 2 and cluster 3 neurons reveals that the fluctuations in both clusters are largely positively correlated, and that the fluctuations in activity of cluster 2 neurons at time *t* predict the fluctuations in activity of cluster 3 neurons for times *t*^′^ > *t* (*p* < 10^−8^, permutation test; Fig. 6D). The observations suggest that the time constant of integration increases with ascending cluster number (or that the degree of integration *leak* decreases with ascending cluster number).

### Representation of momentary motion evidence by T_in_ neurons

The transient effect of the motion pulses on the activity of cluster 1 neurons made us wonder whether these neurons resemble other neurons in LIP that mimic responses of direction selective neurons in area MT (Freedman and Assad, 2006). The Steinemann, Stine et al. (2024) dataset also contains such neurons, termed M_in_—for motion in response field—neurons. Fig. S7A shows firing rate averages from leftward and rightward preferring M_in_ neurons in the random dot motion task. For the M_in_ neurons, the firing rate traces associated with different motion coherences are predominantly parallel. They do not resemble the ramp-like dynamics associated with evidence accumulation. Some M_in_ neurons show selectivity for contraversive motion 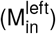 and others for ipsiversive motion 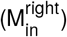(Fig. S7A–C, top and bottom panels respectively).

In contrast to T_in_ neurons, M_in_ neurons do not show persistent activity in the memory-guided saccade task (Fig. S7B) but show direction selectivity in the passive motion viewing task (Fig. S7C). However, during the random dot motion task, M_in_ and 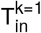 neurons display similar responses in that they are direction selective but represent neither evidence accumulation nor decision termination. Note the similarity of the traces between the M_in_ neurons (Fig. S7A) and the 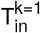 neurons (Fig. 4D, *top*).

We wondered if the 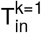 neurons would share signals with the M_in_ neurons, despite the different locations of their response fields. We tested this idea using a noise correlation analysis similar to that shown in Fig. 6B-D, but where the correlations are calculated between 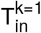 and M_in_ neurons. That is,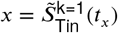 and 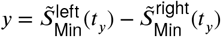. As before, the tilde in these expressions indicates the use of standardized residual values, for each motion coherence. The heat map (Fig. S7D) illustrates the correlation of these residuals across trials. Correlations are stronger for off-diagonal elements where the activity of the cluster 1 neurons lags the activity of the M_in_ neurons by approximately 100 ms. Correlations are significantly higher for *t*_*x*_ > *t*_*y*_ than for *t*_*y*_ > *t*_*x*_ (*p* < 10^−8^; permutation test). That is, fluctuations in the activity of M_in_ neurons predict changes in 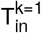 neurons at later times. The finding is consistent with the idea that M_in_ neurons drive 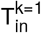 neurons, although a common information source with different latencies cannot be excluded.

### Information about choice and accuracy evolves over time

So far we have mainly focused our analyses on a time window just before the choice report. Here we look for choice and accuracy signals outside the presaccadic window to characterize the time course of information about choice and accuracy. We construct two population signals by projecting the neural activity in the coding directions defined by ***β***_**choice**_ and ***β***_**conf**_. Neural activity is obtained by binning the spike counts of T_in_ neurons in sliding windows of 100 ms. At each time *t*, we compute the area under the ROC curve (AUC) obtained from the projections onto the choice and accuracy coding directions. The AUC values indicate how well the projections discriminate between left and right choices and between correct and incorrect choices, respectively.

Both AUC values peak near the time of reporting (Fig. S8). Unsurprisingly, the decoding of choice is more veridical than the decoding of choice accuracy (i.e., the choice predictions better distinguishes left from right choices than the accuracy predictor distinguishes correct from errors). We assessed whether accuracy information lags behind choice, using a latency analysis based on fitting a bilinear “dogleg” function (Lorteije, Zylberberg et al., 2015) to the time course of AUC values. Information about choice diverges from baseline at 0.165 ± 0.02*s* from motion onset; for choice the divergence occurs 0.187 ± 0.04*s* from motion onset (Fig. S8). This difference is not significantly different from zero (in a bootstrap analysis, choice lags confidence in 24.8% of samples). This suggests that information about choice and accuracy are practically contemporaneous.

### The presaccadic confidence signal accommodates an informative prior

In the random dot motion task, confidence is influenced not only by reaction time and motion strength, but also by the prior probability (base rate) of the different response alternatives (Zylberberg et al., 2018). We ask whether the neural representation of choice accuracy that we identified is also sensitive to manipulations of prior probability. We reanalyzed data from Hanks et al. (2011) in which the prior probability that the motion is rightward or leftward was varied in blocks of ∼ 400 trials.

We decoded choice accuracy using the same approach used for the Neuropixels data. We select trials with a contralateral choice and predict whether the choice is correct or incorrect using the neuronal data recorded on that session, again focusing on the presaccadic window. In the experiment of Hanks et al. (2011), only one T_in_ neuron was recorded per session, hence the predictions are less veridical. Nevertheless, the predicted choice accuracy is greater for trials in which the monkey chose the target with the greater base rate (Fig. 7). This holds for each level of motion strength and for both correct and incorrect decisions (Fig. 7 A & B, respectively), consistent with behavioral observations (Zylberberg et al., 2018). Therefore, the neural representation of choice accuracy is not only informed by motion strength and reaction time (Fig. 3), but also by the prior probability of the chosen option (Fig. 7), thus furthering the idea that the T_in_ neurons support the computation of confidence.

**Figure 7.**
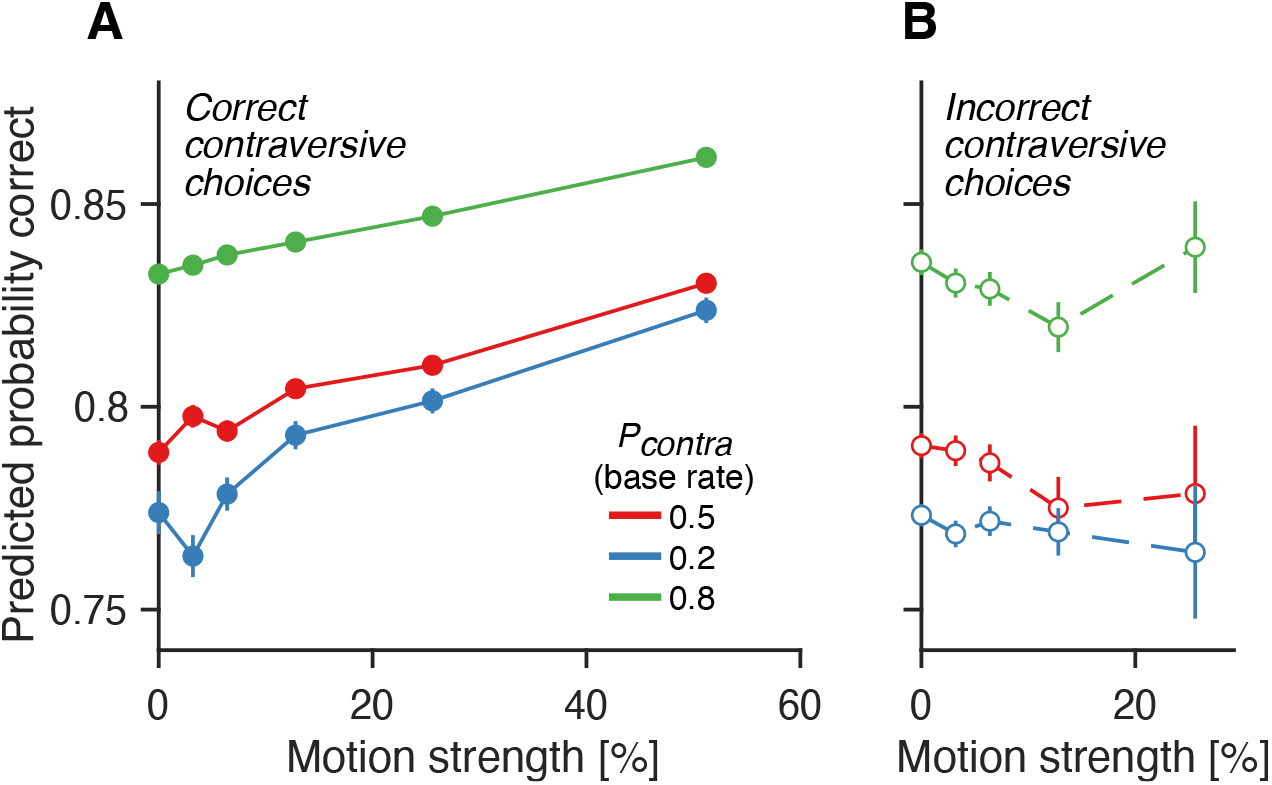
The confidence signal accommodates an informative prior. **(A)** Probability correct inferred from individual LIP neurons from the experiment of Hanks et al. (2011). Monkeys performed blocks of trials in which the prior probability that the target in the neuron’s response field is the one to be rewarded (*p*_contra_) was either 0.5, 0.2 or 0.8. The predicted probability is shown separately for trials of the different blocks (colors). The predicted probability correct increases with motion strength and with the strength of the prior supporting the choice. **(B)** Same as A, but for incorrect choices. In both A and B, the analysis includes only trials in which the monkey chose the target contralateral to the recording site (i.e., the target in the neurons’ response field). Error bars indicate s.e.m. across trials.

## Discussion

### Representation of confidence by T_in_ neurons

Leading models of choice, RT and confidence often assume that confidence depends on the state of the losing race at the moment of choice: the balance of evidence hypothesis (Vickers, 1979; Kiani et al., 2014; Hellmann et al., 2023). This assumption is motivated by a feature of all accumulation models of decision-making— race, attractor and drift-diffusion—that the accumulated evidence for the chosen alternative reaches a positive criterion, or upper bound, at the moment of choice, and therefore cannot provide a graded prediction of accuracy (i.e., confidence). This reasoning turns out to be incorrect. Using high density macaque Neuropixels recordings, we confirm that the population firing rate of T_in_ neurons is indeed stereotyped at the time of decision termination. However the population comprises subsets of neurons with systematic differences in their firing rate dynamics, and this heterogeneity is capable of supporting an estimate of confidence.

This dialectical set of observations is worthy of further dissection. The LIP T_in_ neurons are a functionally defined cell type that exhibits persistent activity associated with saccadic intention and directed attention. The population average firing rate represents the drift-diffusion decision variable that mediates the choice and reaction time on single decisions (Steinemann, Stine et al., 2024), and features of this signal are associated with bursting activity in the superior colliculus that ultimately terminates contraversive decisions (Stine et al., 2023). Yet, not all T_in_ neurons reach a stereotyped level of activity at the end of the decision, and the way each neuron departs from the average is consistent across multiple decisions.

The heterogeneity appears to reflect systematic differences in the dynamical properties of single T_in_ neurons. We arrived at this conclusion indirectly, by first characterizing the response of each T_in_ neuron in the presaccadic window to choice, RT, and motion strength. We clustered the T_in_ neurons based on their encoding of these variables, selecting three clusters. Cluster 1 reflects the motion strength at the end of a contraversive choice; cluster 3 shows a gradual increase in firing over time; cluster 2 is largely independent of motion strength and RT (similar to the average). From the average activity of clusters 1 & 3, we could predict the accuracy of the choice with the same fidelity as when using the full population. The clusters differed in their response to brief motion pulses, in a manner consistent with different time constants of integration (Fig. 6). Such variation might be supported by different levels of recurrence in the neural circuit (Usher and McClelland, 2001; Wong and Wang, 2006; Lange et al., 2021; Zylberberg et al., 2009).

A comparison of the vectors of weights (i.e., the coding direction in state space) that are most informative about choice and accuracy revealed that their cosine similarity is relatively low (Fig. 5D). Therefore, a readout of these two signals from the population of T_in_ neurons would rely on different weightings. However, this dissimilarity does not imply that distinct subsets of T_in_ neurons are dedicated exclusively to choice or confidence. Instead, we favor the interpretation that choice is informed by the activity of all T_in_ neurons. This interpretation is supported by several observations: (i) the regression coefficients used for clustering vary along a continuum (Fig. 4A,D), (ii) T_in_ neurons from different clusters exhibit similar responses in the mapping tasks (memory-guided saccades and passive motion viewing) (Fig. S5), and (iii) the trial-average activity of T_in_ neurons (a single value per trial) contains as much information about choice as all neurons in cluster 2 (Fig. 5G). The misalignment between the weights assigned by the choice and accuracy decoders suggests that neurons that do not encode response time or motion strength contribute to choice decoding but not to accuracy decoding (Fig. 5A,G).

### Behavioral and neural features of confidence

The putative confidence signal reproduces features of confidence reports from humans in tasks similar to the one performed by the monkeys. These features include the relationship between confidence and decision difficulty (as determined by motion strength), prior probability, accuracy, and reaction times (Kiani et al., 2014; Zylberberg et al., 2018). A recent study showed that the monkeys’ confidence, ascertained through bets, display similar behavioral features (Vivar Lazo, 2024). Accounting for these features is thought to require knowledge of the decision time and/or the accumulated evidence for the unchosen alternative (Vickers, 1979; Kiani et al., 2014; van Den Berg et al., 2016; Brus et al., 2021; Smith and Vickers, 1988; Zylberberg et al., 2018, 2012; Hellmann et al., 2023; Moreno-Bote, 2010; Rolls et al., 2010; Wei and Wang, 2015). However, all the information needed to account for these behavioral features is contained in the activity of the T_in_ neurons at the end of the decision.

The presaccadic confidence signal we identified may reveal other features of confidence not explored here, including the positive evidence bias (PEB)—the tendency for confidence to rely more on sensory evidence favoring the chosen option (positive evidence; PE) than the non-chosen option (negative evidence; NE) (Zylberberg et al., 2012; Aitchison et al., 2015; Peters et al., 2017; Maniscalco et al., 2016; Mazor et al., 2023; Samaha and Denison, 2020; Sepulveda et al., 2020; Vivar Lazo, 2024; Mazor et al., 2023). The PEB can be explained by models in which the winning race (i) assigns greater weight to PE than NE and (ii) contributes more strongly to confidence than the losing race (Zylberberg et al., 2012; Paz et al., 2016; Maniscalco et al., 2021). The present study provides support for the latter requirement. Whether the putative confidence signal we identified is more sensitive to PE than NE can be tested in tasks where PE and NE can be manipulated independently (e.g., Lorteije, Zylberberg et al., 2015; Brunton et al., 2013). Other features of confidence that need to be explored in suitable tasks include the increase in confidence with response time when stimulus duration is controlled by the experimenter (Irwin et al., 1956; Vickers et al., 1985; Kiani and Shadlen, 2009; Zylberberg et al., 2016), under time pressure (Vickers and Packer, 1982), and when the evidence is more stochastic (Zylberberg et al., 2014, 2016).

Neurophysiological studies in non-human animals have identified neural correlates of confidence in several brain areas, including the superior colliculus (Odegaard et al., 2018), pulvinar (Komura et al., 2013), orbitofrontal cortex (Kepecs et al., 2008; Lak et al., 2014; Masset et al., 2020), LIP (Kiani and Shadlen, 2009; Vivar Lazo, 2024), supplementary eye fields (So and Stuphorn, 2016; Middlebrooks and Sommer, 2012), visual cortex (Fetsch et al., 2014; Zylberberg et al., 2016; Boundy-Singer et al., 2024), and the midbrain (Lak et al., 2017). Kiani and Shadlen (2009) examined a version of the random dot motion task where the experimenter controlled the stimulus duration. Monkeys had the option to opt out of making decisions to receive a small but guaranteed reward. Kiani & Shadlen suggested that the monkey’s decision to choose or waive the sure bet is mediated by the computation of confidence which depends on (*i*) the state of the decision variable at the end of the evidence stream, and (*ii*) the random dot motion duration from onset to offset. They proposed that LIP contributes to this computation by representing the decision variable through the average activity of T_in_ neurons. Building on this, we show that variation in how individual T_in_ encode motion strength, response time and prior probability enables a linear readout of expected accuracy, suggesting that the representation of confidence by LIP neurons may be more explicit than previously appreciated (Pouget et al., 2016).

### Limitations of the study

A key contribution of our study is the realization that confidence in a decision can be estimated at decision termination simply as a weighted average of the activity of LIP neurons whose response fields overlap with the chosen target. That said, the surprising fact that this information is present does not imply that the monkey exploits it. Additional experiments would be needed to evaluate such an assertion (Ritchie et al., 2019). In our case, this concern is mitigated by three observations: (*i*) we used simple, linear decoders, which implies that it is easy for downstream areas to read confidence from the activity of T_in_ neurons, (*ii*) validated the prediction against many behavioral features of confidence, and (*iii*) focused on a specific moment at the end of the deliberation when these neurons are presumably communicating important information (e.g., where to move the eyes) to downstream areas.

An obvious limitation to the study is the absence of a behavioral assay for the monkey’s confidence, thus precluding a direct comparison between neural activity and behavioral confidence reports within individual animals. Confidence reports cannot be directly collected from nonhuman animals and must instead be inferred through indirect measures, such as betting behavior, willingness to wait for a reward, or opting out of a choice. An advantage of our design is that it demonstrates the presence of neural signals sufficient to estimate confidence, even in monkeys that were never explicitly trained to report it or use it for subsequent decisions (Lebreton et al., 2015). This also ensures that the observed neural representations are not confounded by the behavioral planning required to report confidence. However, our design precludes direct comparisons between neural activity and behavioral confidence reports within individual animals. With access to a behavioral measure of the monkeys’ confidence, we could have performed a decoding analysis similar to the one conducted here, but with confidence as the dependent variable instead of accuracy. We would predict such an analysis to yield results consistent with our findings, although any discrepancies might also be informative. For example, if the neural state-space direction most informative about confidence differs from the direction most informative about accuracy (e.g., cosine similarity significantly less than 1), this might explain why confidence often appears to make suboptimal use of the information available for constructing the choice (Rahnev and Denison, 2018).

Another limitation of our paradigm is its inability to disentangle confidence in making the correct decision from the expectation of obtaining a reward. This ambiguity applies to most neurophysiological studies of confidence. Resolving this distinction may require tasks where reward magnitude is manipulated independently of other variables (Rorie et al., 2010; Fan et al., 2024, 2020; Doi et al., 2020; Fan et al., 2018). Fan et al. (2024) employed such a task and found that both accuracy and reward expectation are represented in the caudate nucleus of the basal ganglia and the frontal eye fields. High-density population recordings in these tasks may help clarify whether and how the heterogeneity of T_in_ neurons supports the estimation of accuracy or reward expectation, or both.

Our study also underscores the importance of characterizing single-neuron activity, even when conducting population-level analyses. The heterogeneity of neuronal responses was observed within a small population of LIP neurons (T_in_) that share the property of displaying persistent activity toward the contralateral choice target. This property was identified in a mapping task separate from the main task. Without this characterization, the heterogeneity we identified would not bear on the predictions of the balance of evidence hypothesis. Therefore, single-neuron and population-level analyses might best be conceived as complementary rather than competing views (cf., Barack and Krakauer, 2021).

## STAR Methods

### Key resources table

**Table 1.** Resources and their identifiers.

| RESOURCE CATEGORY | SOURCE | IDENTIFIER |
| --- | --- | --- |
| **Experimental data** |  |  |
| Neural & behavioral | Steinemann, Stine et al. (2024) | https://doi.org/10.5281/zenodo.7946011 |
| Neural & behavioral | Hanks et al. (2011) | Not publicly available |
| Behavioral | van Den Berg et al. (2016) | Not publicly available |
| Behavioral | Zylberberg et al. (2018) | https://github.com/arielzylberberg/ZylberbergWolpertShadlen_Neuron2018 |
| **Software and algorithms** |  |  |
| Original code | This paper | TBD |
| MATLAB | MathWorks | https://www.mathworks.com/products/matlab.html |

### Resource availability

#### Lead contact

Further information and requests for resources should be directed to and will be fulfilled by the lead contact, Ariel Zylberberg.

## Data and code availability

This study did not generate new data. All original code has been deposited at Github and is publicly available as of the date of publication.

Any additional information required to reanalyze the data reported in this paper is available from the lead contact upon request.

## Materials availability

This study did not generate new unique reagents.

## Experimental model and study participant details

This study did not generate new data.

## Methods details

### Behavioral tasks

#### Random dot motion task

In the main task, the monkeys had to decide the net direction (leftward or rightward) of a circular patch of limited-lifetime, dynamic random dots and report their choice when ready by making a saccadic eye movement from the central fixation to the left or right choice target. The monkey initiates at trial by directing the gaze to a central fixation point. After 0.25–0.7 s (truncated exponential with time constant *λ*=0.15 s), two red choice targets (diameter 1 dva; degrees of visual angle) are presented in the left and right visual fields. After a random delay (0.25-0.7 s, *λ*=0.4 s), the random dot motion stimulus is displayed until the monkey initiates a saccadic eye movement to report its choice.

The random dot motion comprises limited lifetime dots displayed within a circular area (diameter 5 dva) centered on the fixation point. The dot density is 16.7 dots · dva^−2^· s^−1^. The direction and strength of the motion are chosen pseudorandomly on each trial, such that the coherence, *C ∈* {±0%, ±3.2%, ±6.4%, ±12.6%, ±25.6%, ±51.2%}. The sign of *C* determines the direction of motion; positive values indicate leftward motion. For *C* = 0%, the sign indicates the random direction that is rewarded on that trial. The absolute value | *C* | establishes the motion strength: the probability that a dot displayed in video frame *n* is displaced by Δ*x* in frame *n* + 3 (i.e., 40 ms later). Otherwise the dot is replaced by a new dot at a random position. The displacement, Δ*x* = ±0.2 dva, is consistent with apparent motion speed of 5 dva per s (see Roitman and Shadlen, 2002, for further details). Monkeys are rewarded for making a saccadic eye movement to the correct choice target. On trials with 0% motion coherence, either saccadic choice is rewarded with a probability of 0.5. Incorrect responses are penalized by increasing the inter-trial interval by up to 3 seconds (see Stine et al., 2023, for further details). On approximately half of the trials, the motion coherence is incremented or decremented for 100 ms by 4% coherence for monkey M and 3.2% for monkey J (Stine et al., 2023). The onset time of the *pulse* is chosen randomly from a truncated exponential distribution: *t*_*min*_ =0.1 s to *t*_*max*_ =0.8 s from motion onset (*λ* = 0.4 s). Monkey M completed 9,684 trials across five sessions, while Monkey J completed 8,142 trials in three sessions.

#### Control tasks

Monkeys also completed two additional tasks in each session: a passive motion viewing task and a memory-guided saccade task. In the passive motion viewing task, the monkey views ±51.2% coherent motion for 0.5 s (and for 1 s on a small number of trials in one session). The task matches the main task but without choice targets. The monkey is rewarded for maintaining fixation during the motion presentation.

In the memory-guided saccade task (Hikosaka and Wurtz, 1983; Gnadt and Andersen, 1988), a target was briefly flashed (200 ms) at a pseudo-random location in the visual field. After a variable delay (0.2-0.9 seconds for monkey M, *λ* = 0.3 seconds; 0.3-1.3 seconds for monkey J, *λ* = 0.2 seconds), the fixation point was extinguished and the monkey had to make a saccadic eye movement to the remembered location of the target. The monkey was rewarded if the gaze was within ±2.5 degrees of visual angle of the target location.

#### Biased prior probability task

The analysis of prior probabilities makes use of previously published single neuron recordings from two other monkeys that performed the same task as Steinemann, Stine et al. (2024). However, Hanks et al. (2011) varied the prior probability that the rewarded choice was left or right. In blocks of trials, the sign of the motion coherence was biased in favor of positive or negative. They also included blocks with a neutral (i.e., uninformative) prior. In blocks with neutral priors, both targets had an equal 50% chance of being correct. In biased conditions, one direction had an 80% probability of being correct and the other had a 20% probability of being correct, except for a small number of sessions in one monkey where a 67-33% prior was used (these data were not included in our study nor in Hanks et al. 2011).

Sessions typically began with a block of 200-400 trials under a neutral prior. The monkeys then completed 300-600 trials in which the prior favored one of the targets, with the most-likely target being the one chosen least often during the neutral prior block. To signal to the monkeys which target was more likely, each biased block was preceded by 20 trials of 100% coherent motion toward the more likely target. These trials were not included in our analysis. In some sessions, monkeys completed an additional block with a prior favoring the opposite target. See Hanks et al. (2011) for details.

#### Combined choice-confidence task in humans

van Den Berg et al. (2016) asked participants to discriminate the direction of motion of a random dot motion display similar to that of Steinemann, Stine et al. (2024) and Hanks et al. (2011). Subjects held the handle of a vBOT manipulandum used to record the position of the handle at 1,000 Hz (Howard et al., 2009). A horizontal mirror projecting a downward facing CRT monitor prevented subjects from seeing their arm. A chin rest ensured that the viewing distance was approximately 40 cm.

Participants reported choice and confidence simultaneously by moving the handle to one of four circular targets displayed at the corners of a 17 cm x 17 cm square. The two targets on the left corresponded to a leftward motion choice, and the two on the right corresponded to a rightward motion choice. In half of the blocks, the two top targets corresponded to a high-confidence choice and the bottom targets to a low-confidence choice; in the other half, the mapping was reversed such that the bottom targets corresponded to high-confidence and the top targets to high-confidence. To motivate participants to make calibrated confidence reports, the high- and low-confidence targets had different payoffs for correct and incorrect choices. The low confidence targets gave a 1 point reward for a correct choice and a 1 point loss for an incorrect choice. The high-confidence target gave 2 points for a correct choice and a loss of 3 points for an incorrect choice.

Fig. 3A reproduces data from a representative participant (Subject 2 in Figure 2 of van Den Berg et al. 2016), who completed 9,023 trials over 12 experimental sessions. Data from the other participants is shown in Fig. S1. Details of the experimental procedure should be sought in the original publication (van Den Berg et al., 2016).

### Neurophysiological recordings (LIP)

#### Main task

Steinemann, Stine et al. (2024) used a prototype “alpha” version of Neuropixels1.0-NHP45 probes (developed by IMEC and HHMI-Janelia) to record multiple single-unit activities in the ventral part of area LIP (LIPv). Steinemann, Stine et al. (2024) used anatomical MRI to localize LIPv and used single-neuron recordings (Thomas Recording GmbH) to verify that the activity conformed to known physiological properties of LIPv before proceeding with multi-neuron recordings. The Neuropixels probes are equipped to record from 384 of the 4,416 available electrical contacts distributed along their 45 mm long shaft. Data was only recorded from the 384 contacts closest to the probe tip (Bank 0), covering 3.84 mm. The reference and ground signals were directly connected to each other and to the monkey’s headpost. A total of 1,084 neurons were recorded over eight sessions, with each session yielding between 54 and 203 neurons (see Table 2 of Steinemann, Stine et al. 2024 for details).

#### Biased prior probability task

Hanks et al. (2011) recorded fifty-two neurons from the LIP area of two rhesus monkeys. Recordings were made using standard methods for extracellular recording of action potentials from single neurons (Roitman and Shadlen, 2002). See Hanks et al. (2011) for details.

### Quantification and statistical analysis

#### Race model of decision making

In the race model, two drift-diffusion processes, *x*_*L*_(*t*) and *x*_*R*_(*t*), compete until one of them reaches an upper bound. The first to reach the upper bound determines choice and RT. *x*_*R*_ accumulates evidence for right minus left, and *x*_*L*_ accumulates evidence for left minus right. The dynamics of the decision variables is described by the following difference equations:

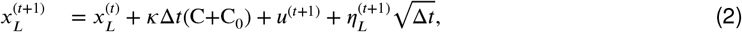

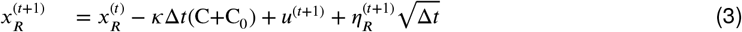

where *κ* is a measure of the signal-to-noise, Δ*t* = 0.005s is the time step, C is the motion coherence (positive for leftward motion), C_0_ is a bias term and *η* is zero-mean normally distributed noise with unit variance. *η*_*L*_(*t*) and *η*_*R*_(*t*) are sampled from a bivariate Normal distribution such that the correlation between them is *ρ* = −0.7. At time *t* = 0, *x*_*L*_ = *x*_*R*_ = 0.

The urgency signal *u*^(*t*+1)^ decreases the amount of evidence needed to trigger a response as time progresses (Hanks et al., 2011). For times *t*<*d*, the value of *u*^(*t*+1)^ is zero, indicating no urgency. For times greater than *d, u*^(*t*+1)^ assumes a constant value equal to *a*Δ*t*, where *a* is a parameter that represents the linear rate of rise of the urgency signal.

The decision variables in the model cannot drop below a lower reflective bound. If an update to a decision variable would result in a value lower than *B*_ref lect_, the variable’s value is set to *B*_ref lect_ for that time step. That is:

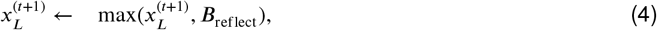

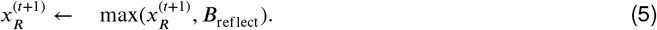

The lower, non-absorbing bound simply instantiates the fact that firing rates must be ≥ 0.

The decision terminates when one of the races reach an upper bound at *B*. The choice is leftward (rightward) if *x*_*L*_ (*x*_*R*_) reaches the bound first. The decision time is the time taken to reach the bound. The reaction time is the sum of the decision time and a non-decision time, which is normally distributed with a mean of *μ*_*nd*_ and standard deviation σ_*nd*_.

The free parameters of the model are Θ = {*κ, B, a, d*, C_0_, *μ*_*nd*_, σ_*nd*_}. We use simulations to fit them to data. Data from each monkey were fit separately. For a given set of parameters, we simulated 10 times as many trials as were completed by the monkey. For each combination of choice and motion coherence, we fit the distribution of decision times obtained from the model with an Epanechnikov (parabolic) kernel to obtain a smooth probability density function of decision times. The distribution of decision times is convolved with the distribution of non-decision times to obtain a probability density function of reaction times. We compute a separate p.d.f. of RTs for each combination of choice and motion coherence, and use them to calculate the likelihood of the parameters given the single-trial choice and reaction time data. We use BADS (Acerbi and Ma, 2017) to find the maximum-likelihood parameters. The best-fitting parameters are shown in Table S1.

### Data analysis

#### Preprocessing of neuronal data

Our study focuses on T_in_ neurons, i.e., those that show persistent activity during saccade planning to the target contralateral to the recording site. For the Neuropixels data, T_in_ neurons were identified post hoc using a memory-guided saccade task (Steinemann, Stine et al., 2024; Stine et al., 2023). Hanks et al. (2011) used the same task to identify neurons with spatially selective persistent activity and to place targets within the response field of these neurons.

Unless otherwise stated, neurophysiological analyses are based on the number of spikes emitted by each T_in_ neuron in the 100 ms epoch that ends 50 ms before the initiation of the saccadic eye movement used to report the choice. We refer to this time interval as the presaccadic window.

#### Accuracy decoder

In each of the 8 sessions, we trained two decoders to predict the accuracy of the monkey’s choice, using only contralateral (left) or ipsilateral (right) choices, respectively. We trained the decoder using simple logistic regression:

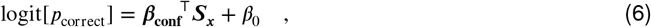

where 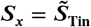 is the standardized spike count of each T_in_ neuron in the interval from 150 ms to 50 ms before saccade onset. Standardization (i.e., z-score) is applied to each neuron independently. The fitted coefficients, ***β***_**conf**_, establish a vector the same size as the number of simultaneously recorded T_in_ neurons. We refer to this vector as a coding direction in neuronal state space. *β*_0_ is a constant that captures the accuracy rate over all stimulus conditions, which is typically much better than chance (typically, *p*_correct_ > 0.7).

We apply the same strategy to decode from subpopulations of neurons by redefining ***S***_***x***_. For example, Fig. 5A shows the result of applying the regression model (Eq. 6) to the subset of T_in_ neurons belonging to each cluster. For the analysis of the data from Hanks et al. (2011), *S*_*x*_ contains the standardized spike counts of the single T_in_ neurons recorded in separate experiments. We use the same presaccadic window as in the analysis of the Neuropixels data.

To derive the confidence estimates, the decoders are trained using 10-fold cross-validation. The data are divided into 10 groups, each containing an approximately equal number of trials selected randomly. One group is used as the prediction set, while the remaining nine groups are used for training. This process is repeated 10 times, ensuring that the confidence estimates for each trial are based on a prediction. For the analyses shown in Fig. S8, we derive ***β***_**choice**_ and ***β***_**conf**_ using all trials instead of using cross-validation. This approach allows us to obtain a single set of regression weights per session, rather than 10 sets as produced by the cross-validation method.

The confidence estimates are used to calculate the area under the ROC curve (AUC). Fig. 2A exemplifies the distributions that are used for the ROC analysis. For the analyses shown in Fig. 2B and Fig. 5E, AUC values were calculated separately for each session. The statistical comparisons use a one-tailed paired *t-test* applied to the logit-transformed AUC values from each session. Elsewhere, AUCs are not computed per session, but rather the confidence estimates from all sessions are pooled before calculating a single AUC. Standard errors were computed using bootstrapping (*N* = 5, 000 samples).

We also use bootstrap samples to determine if there is a significant difference between two AUC values. For instance, to determine if the AUC_conf_ obtained using only the cluster 1 neurons is larger than that obtained using only the cluster 2 neurons (Fig. 5A), we bootstrap to obtain *N* = 5, 000 AUC_conf_ values for each cluster. We then compare the AUC values for all *N* = 25 × 10^6^ pairwise comparisons and determine significance as the proportion of comparisons for which the AUC value from cluster 1 neurons is larger than that from cluster 2 neurons.

#### Choice decoder

We use the same spike-count standardization (z-scoring), analysis time window, and cross-validation method to predict the monkey’s choice on each trial:

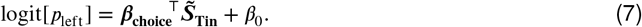

Here, *β*_0_ reflects the monkeys bias for or against a left choice, not the monkey’s accuracy. Unlike the accuracy decoder, the choice decoder is trained on trials with both contralateral and ipsilateral choices.

#### Reaction time decoder

We also use a logistic decoder to assess whether T_in_ neurons contain, just before the response, information about reaction time. The model is:

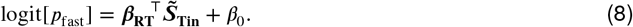

It was fit independently for each session using only trials with contraversive (left) choices. Fast and slow responses were defined relative to the median RT. We use the same cross-validation method that we used for the accuracy and choice decoders.

#### Latency analysis

We used the cumulative sum (CUSUM) method to determine the latency of direction-selective responses in T_in_ neurons (Ellaway, 1978). Receiver operating characteristic (ROC) analysis was used to assess the directional selectivity of each T_in_ neuron. The area under the curve (AUC) in this analysis represents the degree separation—from 0.5 (complete overlap) to 1 (complete separation)—between the spike count distributions for leftward and rightward motion in single trials. The AUC was calculated from spike counts in the interval 100–400 ms after motion onset. We restricted the analysis to correct trials with reaction times greater than 450 ms and motion coherence greater than 10%. For neurons with an AUC greater than 0.55, we calculated the difference in spike counts (in 25 ms bins) between leftward and rightward choice trials. These differences were then added cumulatively over time, as required by the CUSUM method. Typically, the cumulative difference remains around zero prior to the onset of direction selectivity, and then gradually increases or decreases depending on the neuron’s preferred choice.

To determine the onset of direction selectivity, we fit a “dogleg” function to the cumulative spike sum. This function starts with a flat line from *t*_0_ = 0 and transitions to a linear increase or decrease starting at *t*_1_ > *t*_0_. The end of the flat portion of the fit, which occurs between 0 and 500 ms after the onset of motion, was considered the latency. Using cumulative sums of spikes to estimate latencies helps reduce the effect of neural noise. The fitting step further reduces the influence of the number of trials on latency estimates, providing an advantage over traditional methods such as t-tests in moving windows (e.g., Lorteije, Zylberberg et al., 2015). The significance of the difference in latency between neurons from different clusters was assessed with a two-tailed *t-test*.

We conducted a similar analysis to estimate the latencies depicted in Fig. S8. We construct bootstrap samples combining the time course of AUC values from individual sessions (N = 5,000 bootstrap samples). A dogleg function was fit to each bootstrap sample, resulting in 5,000 latency estimates. The arrows in Fig. S8 identify the mean latency across the bootstrap samples. The p-value we report is the proportion of bootstrap samples for which the choice signal deviates from baseline later than the confidence signal.

#### Clustering

We use linear regression to explain the spike counts of each T_in_ neuron in the presaccadic window. As independent variables, we used the motion coherence (*C*), the choice and reaction time (RT), and an offset:

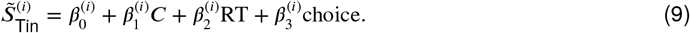

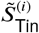 is the standardized spike count of neuron *i* in the aforementioned time interval. The regression analysis was performed separately for each T_in_ neuron, including correct trials only.

We then apply *k-means* clustering, using 3 clusters, to the regression coefficients associated with motion coherence, RT and choice. The cluster labels (1–3) were chosen so that the average of *β*_1_ is smallest for cluster 1 neurons, intermediate for cluster 2 and largest for cluster 3. We confirmed the robustness of the neuron cluster assignments by independently deriving them using either the odd or even trials (Fig. S3). We also fit a regression model similar to Eq. 9, but without the choice and RT terms and considering only correct contralateral choices. The regression coefficients associated with motion coherence are shown in Fig. 4D.

#### Motion pulses

We estimate the time course of the effect of the brief motion pulses on neuronal activity by aligning the spike times of each T_in_ neuron to the onset of the motion pulse. For each time −60<*t*<800 ms relative to the onset of the motion pulse, we calculate the number of spikes emitted in the time epoch between *t* and (*t*+100) ms. Time *t* is advanced in steps of 20 ms. *R*_*i*_(*t, j*) contains the spike counts for neuron *i* emitted at time *t* from pulse onset, in trial *j*. Only trials where the reaction time is at least 150 ms greater than *t*, and those with a motion pulse 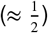 are included in the analysis. We then standardize the spike counts independently for each motion coherence, to obtain 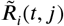. We average 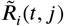 across trials with a leftward (contraversive) pulse, and trials with a rightward (ipsiversive) pulse, and calculate the difference, left minus right, between these averages, to obtain 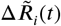.

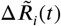 approximates for each T_in_ neuron *i* and time *t*, the influence of the motion pulse on neuronal activity. We average 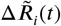 across the *N*_*K*_ neurons belonging to the same cluster *K*,

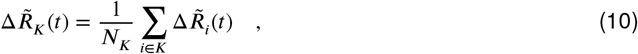

and normalize the average subtracting a baseline,

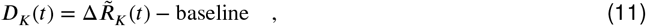

where baseline is the average of 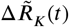 for times *t* between 0 and 0.2*s*. Fig. 6A shows *D*_*K*_ (*t*) for the three clusters.

We use a curve-fitting approach to estimate the rate at which the effect of the motion pulse dissipates over time. We fit *D*_*K*_ (*t*) using a function *f* (*x*) constructed on the following two assumptions: (*i*) the onset latency of the effect of the motion pulse on neuronal activity follows a Normal distribution, (*ii*) the effect dissipates exponentially. Given these assumption, the differential equation describing the time course of *f* (*t*) is

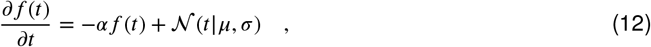

where *μ* and σ are the mean and standard deviation of the Normal distribution (𝒩), and *α* is the reciprocal of the time constant of the dissipation. The solution to this equation is:

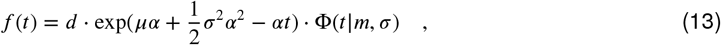

where *d* is a scaling parameter, and Φ(· | *m*, σ) is the cumulative Gaussian distribution with mean *m* = *μ*+σ^2^*α* and standard deviation σ. The fit function has four parameters: {*d, μ*, σ, *α*}. We fit the parameters to minimize the sum over time points of the squared differences between *f* (*t*) and *D*_*K*_ (*t*).

We compared the best-fitting dissipation parameter (*α*) across clusters. We generated bootstrap samples (N=5,000) by selecting with replacement from the pool of neurons that belong to a given cluster. For each of these samples, we compute the parameter *α*. We evaluate the significance of the difference in *α* values in the data by the proportion of bootstrap samples that result in a difference in *α* values as extreme as the one we observed in the data.

#### Cross-correlation analysis

The analysis depicted in Fig. 6B–D is based on the spike counts of neurons from clusters 1–3. Spike counts were calculated in 25 ms windows, aligned to motion onset, up to 50 ms before the reaction time. We compute standardized residuals separately for each time bin, motion coherence and session. Standardized residuals were combined across sessions. The processed signals are referred to as 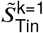,

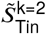 and 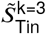. We then calculated the Pearson correlation coefficient between every pair of signals (Fig. 6B–D), for every pair of time steps between 0.2 and 0.8s.

We used permutation tests to asses statistical significance. We define two regions of interest based on the time from stimulus onset in the *x* and *y* dimensions (Fig. 6B). ROI1 is defined by *t*_*x*_ > *t*_*y*_, and ROI2 is defined by *t*_*y*_ > *t*_*x*_, for time time points shown in Fig. 6B. If *y* causally affects *x*, or if *y* and *x* receive a common input but the integration time constant is greater for *y* than for *x*, then the pairwise correlations between *x* and *y* should be greater in ROI1 than in ROI2. We calculated the difference in correlations between two groups, ⟨*ρ*_ROI1_⟩ − ⟨*ρ*_ROI2_⟩, where the expectation is calculated over the time bins within each region of interest (ROI). This difference was then contrasted with those obtained after randomly shuffling the order of trials for one of the dimensions (*N*_shuffles_ = 200). Significance was determined by the probability of achieving a difference as extreme as the one observed in the experimental data.

The same procedure was applied to the cross-correlation analysis shown in Fig. S7D, but with 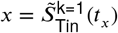 and 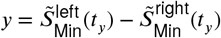, where 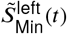 and 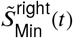 are the standardized residuals obtained from the activity of the M_in_ neurons preferring contraversive and ipsiversive motion, respectively.

## Acknowledgments

We thank Gabriel Stine, Natalie Steinemann, Eric Trautmann, Tim Hanks, Ronald Van den Berg, Kavitha Anandalingam and Daniel Wolpert for sharing the data analyzed in this study.

We also thank Mehdi Sanayei for helpful comments on an earlier version of the manuscript.

## Supplemental information

**Table S1.** Best-fitting parameters of the race model. *ρ* and *B*_rectif_ were not fit but fixed to predefined values.

| | $\kappa$ | $B$ | $a[s^{-1}]$ | $d[s]$ | $\rho$ | $\mu_{nd}[s]$ | $\sigma_{nd}[s]$ | $C_0$ | $B_{rectif}$ |
| --- | --- | --- | --- | --- | --- | --- | --- | --- | --- |
| Monkey M | 14.86 | 1.73 | 1.63 | 0.13 | -0.7 | 0.28 | 0.07 | 0.01 | -1 |
| Monkey J | 12.97 | 0.88 | 0.57 | 0.62 | -0.7 | 0.3 | 0.03 | 0 | -1 |

**Figure S1.**
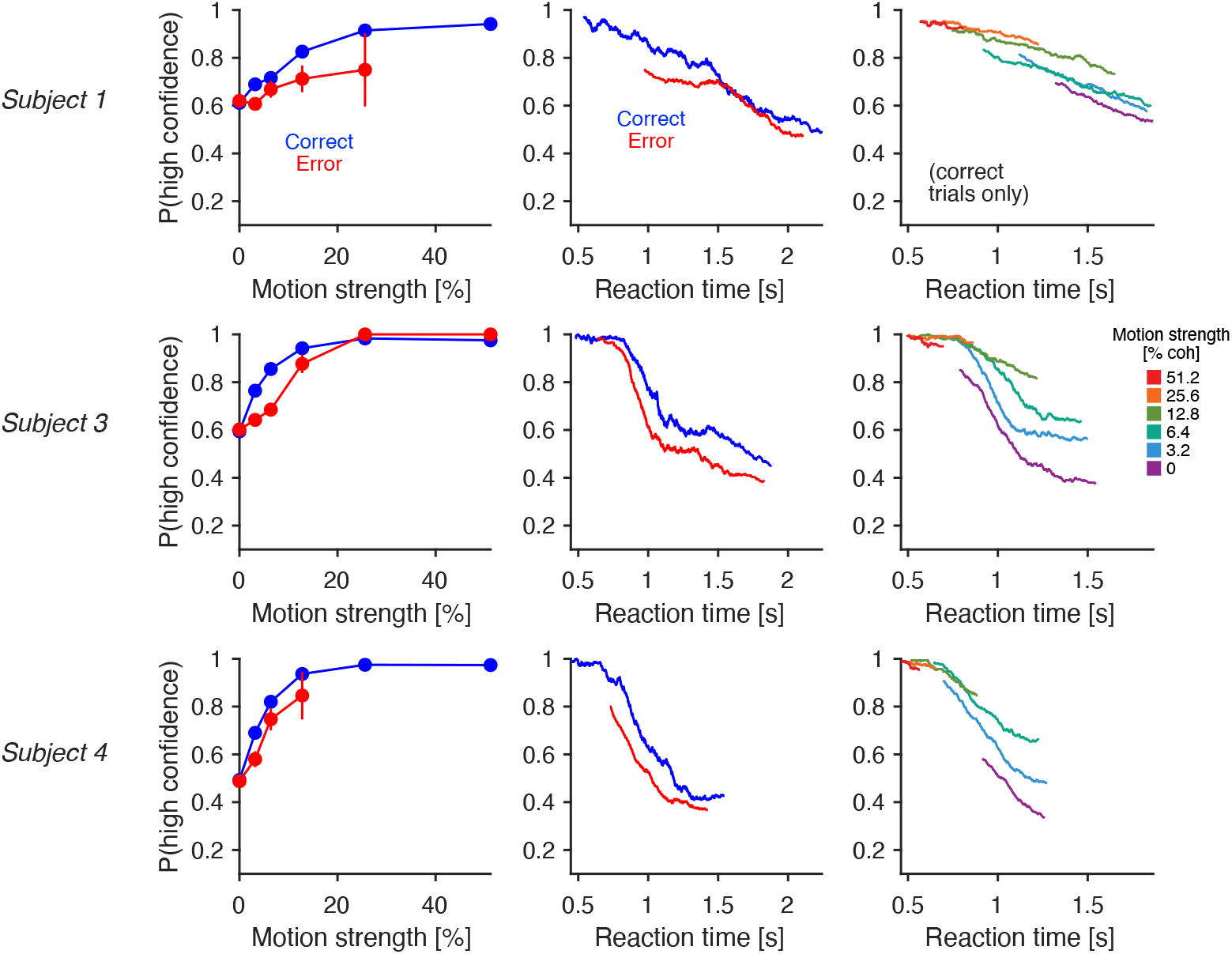
Signatures of confidence in the data of van Den Berg et al. (2016) Same as Fig. 3A for the other three participants in van Den Berg et al. (2016).

**Figure S2.**
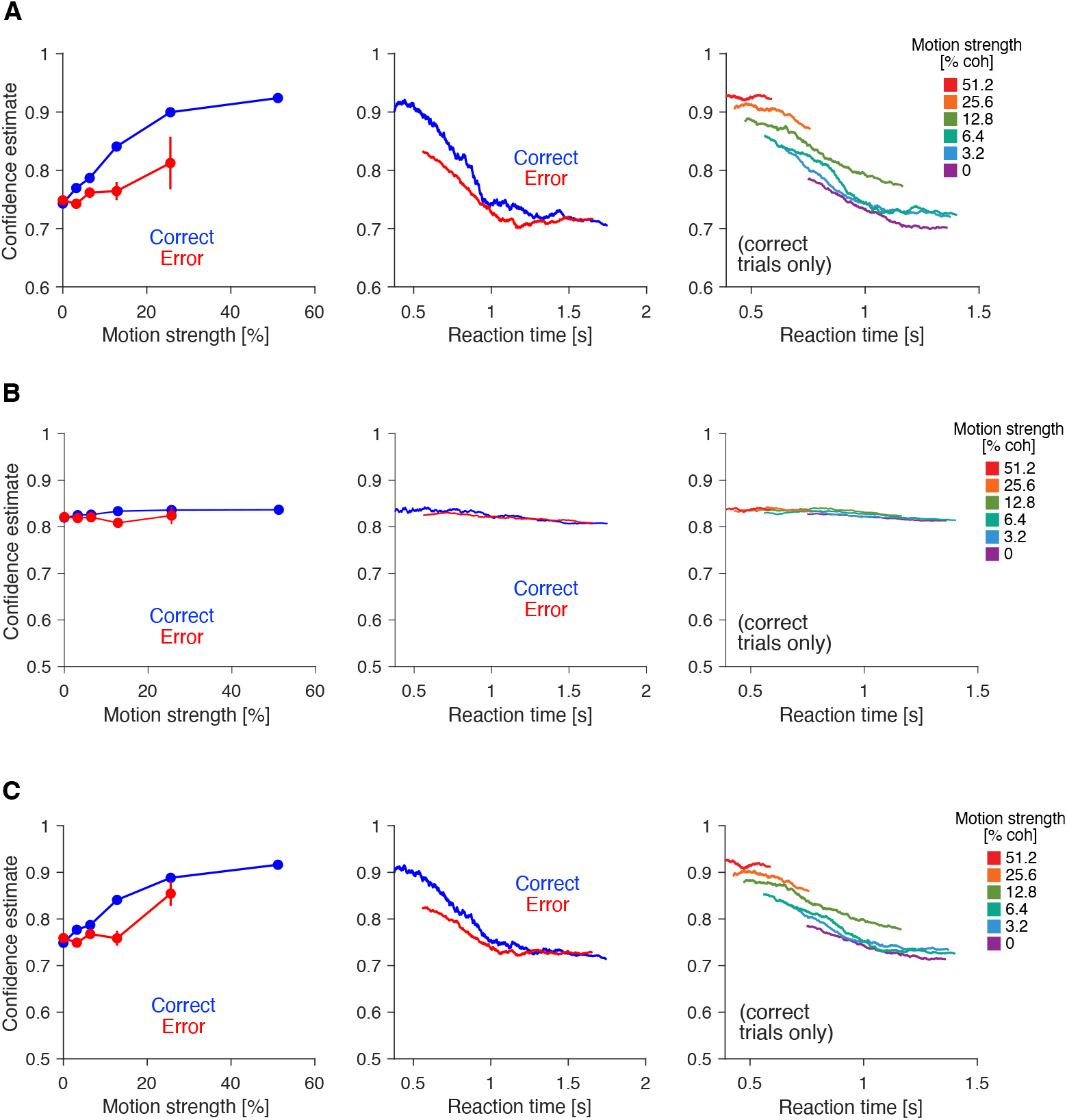
Choice accuracy inferred from neural activity. **(A)** Same as Fig. 3B, except that the confidence estimates obtained from the accuracy decoder are not thresholded into high and low confidence categories. Results are qualitatively similar to those obtained from the behavioral data (Fig. 3A). **(B)** Same as panel A, but for the decoding analysis using gaze data before and after the saccadic eye movement as predictors, instead of neural signals. **(C)** Same as panel A, but using a narrower presaccadic window (from 100 ms to 50 ms before the choice report).

**Figure S3.**
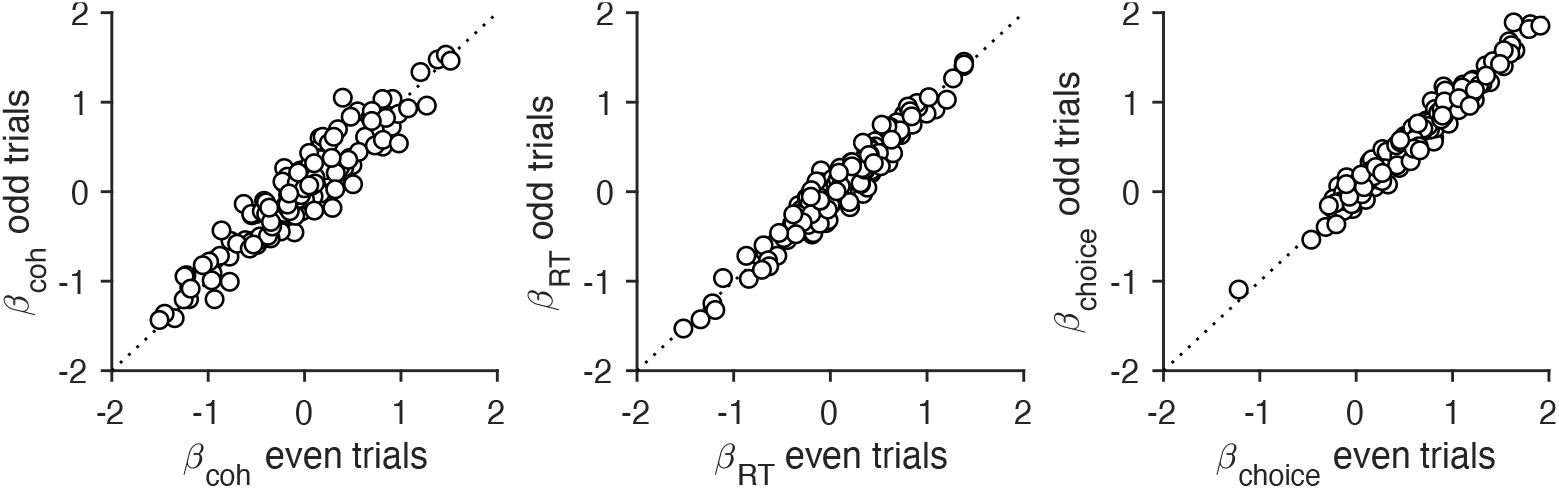
Consistency of the neural representation of motion coherence, reaction time and choice. We use only the odd or even trials to explain the standardized (z-scored) spike counts of each neuron in the presaccadic window as a function of motion coherence, RT, and choice. The figure shows the best-fitting regression coefficients plotted against each other. Each data point corresponds to a different T_in_ neuron. Panels 1-3 correspond to the best-fitting regression coefficients for motion coherence, RT, and choice, respectively. The regression coefficients are highly consistent across independent regression analyses.

**Figure S4.**
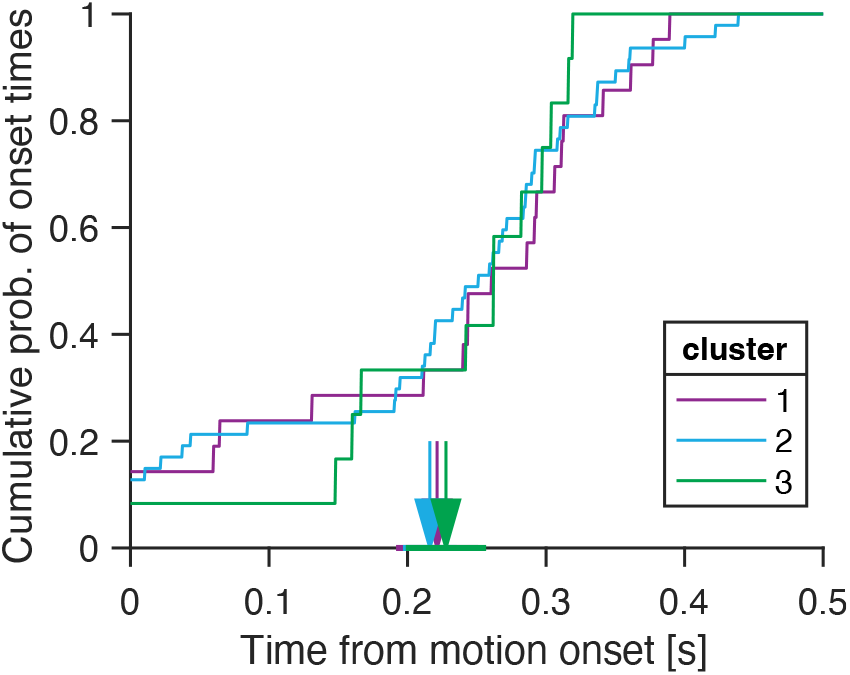
Latency to motion selectivity of individual T_in_ neurons. For each T_in_ neurons separately, we calculate the latency to motion direction selectivity using the CUSUM method (Ellaway, 1978). The cumulative distribution of onset times is shown separately for neurons belonging to the three clusters. The vertical arrows indicate the mean onset time for each cluster, and the horizontal line indicates the standard error of the mean.

**Figure S5.**
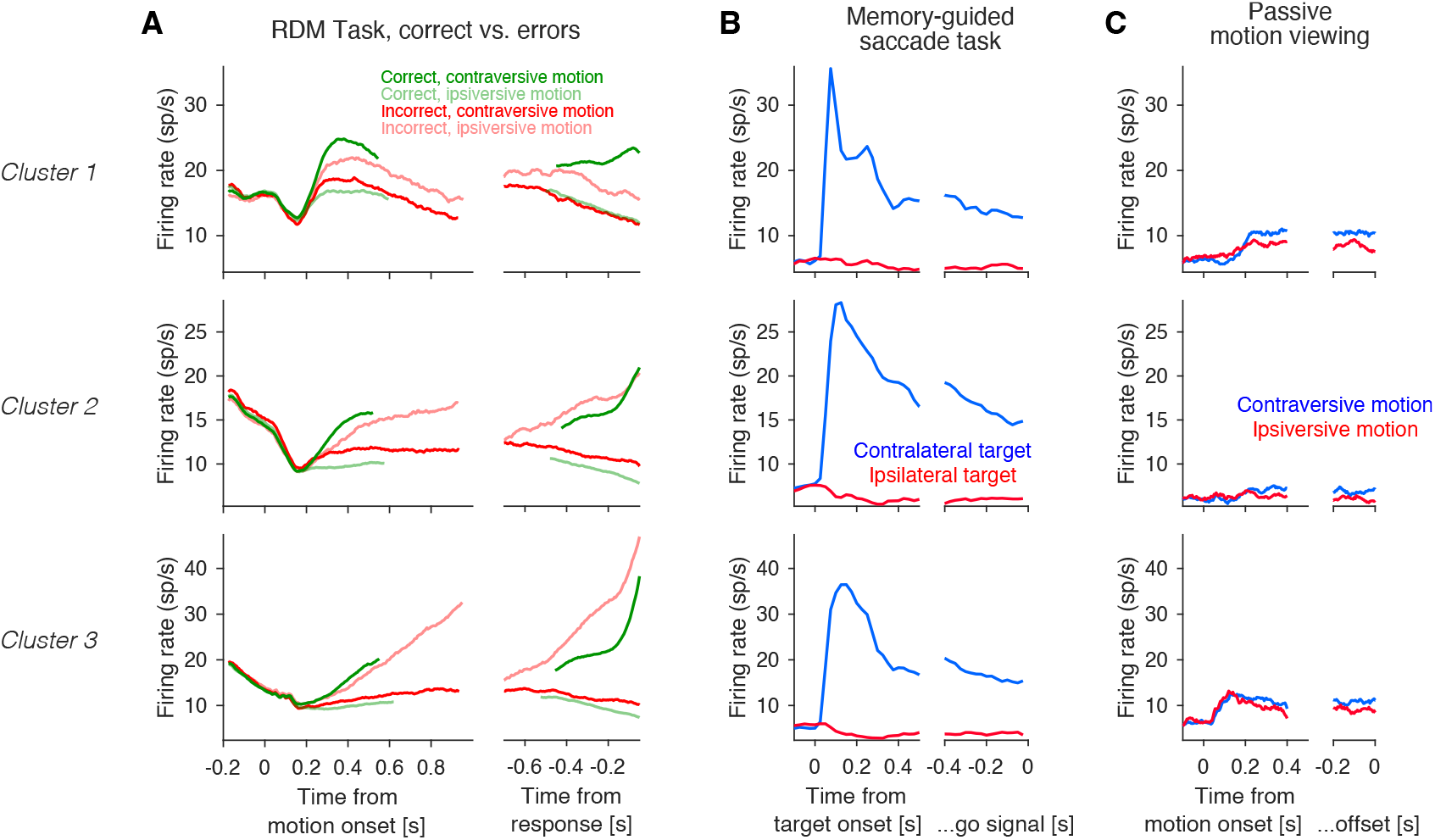
Mean response of neurons from each cluster for correct and incorrect decisions, memory-guided saccades, and passive motion-viewing tasks. **(A)** Random dot motion task. Neurons from all three clusters exhibit stronger responses to contralateral (leftward) choices compared to ipsilateral choices, regardless of whether the choice is correct (green) or incorrect (red). **(B)** Memory-guided saccade task. Blue and red traces correspond to saccades to the target located contralaterally and ipsilaterally, respectively. The horizontal gray bar indicates the time of target presentation. In the panels on the right, neural activity is aligned to the go signal (i.e., the offset of the fixation point). The average time from target onset to the go signal is 0.82 seconds. **(C)** Passive motion–viewing task. Blue and red traces correspond to contraverise (leftward) and ipsiversive (rightward) motion, respectively. Traces are aligned to the onset (left) and offset (right) of the random dot motion stimulus.

**Figure S6.**
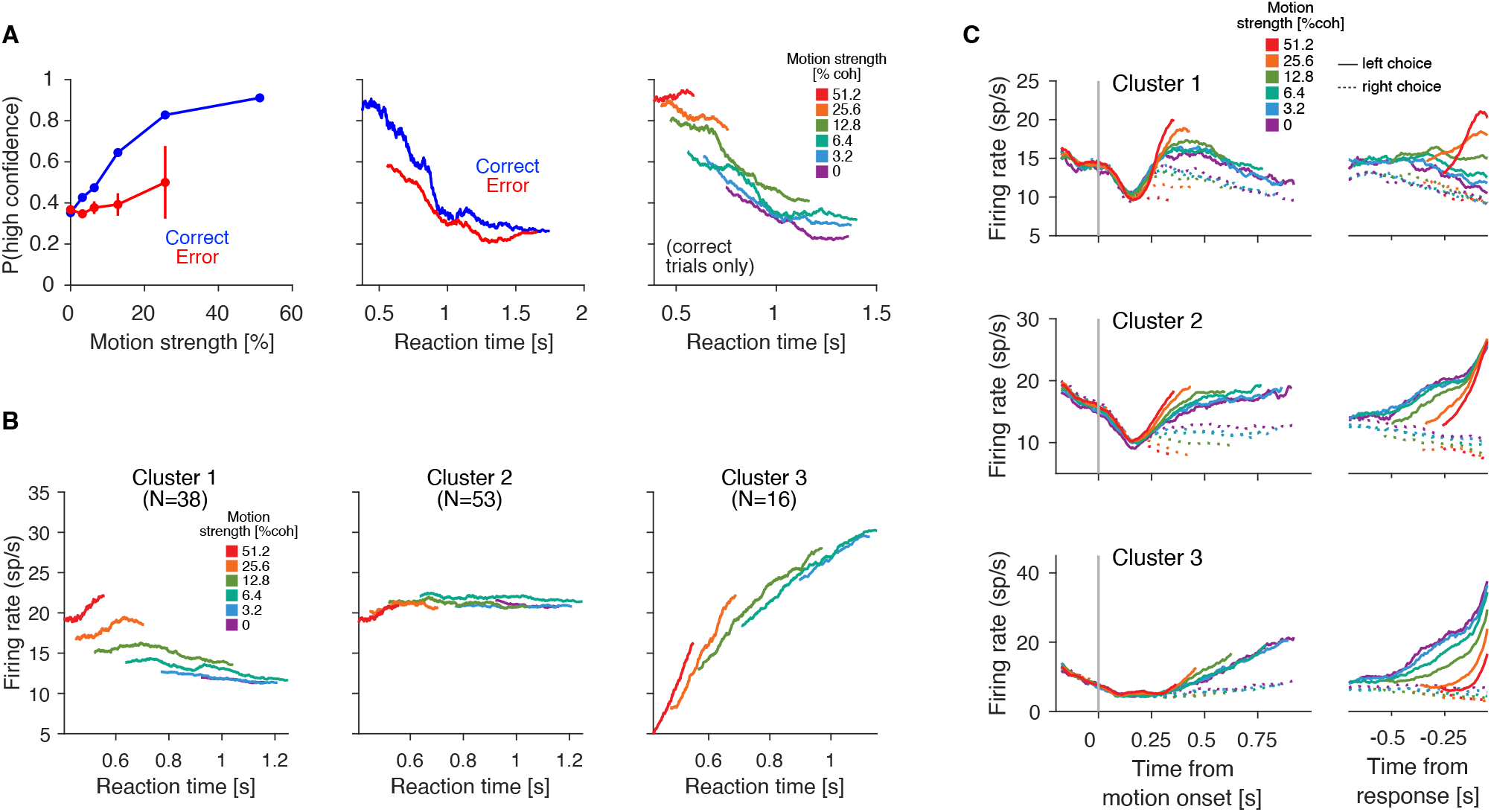
Decoding and clustering without the T_in_ neurons with significant modulation in the passive motion-viewing task. We repeated the decoding and clustering analyses without the T_in_ neurons that significantly discriminated between leftward and rightward motion in the passive motion viewing task. Significance was assessed using a Wilcoxon rank–sum test comparing spike rates on leftward and rightward motion trials. Spike rates were calculated for each trial in the epoch between 0.2s after motion onset and motion offset. Neurons with p-values lower than 0.05 were deemed significant. The results of the decoding and clustering analyses are qualitatively similar to those obtained without excluding these neurons. **(A)** Analysis equivalent to that shown in Fig. 3B. **(B)** Analysis equivalent to that shown in Fig. 4B. **(C)** Analysis equivalent to that shown in Fig. 4C.

**Figure S7.**
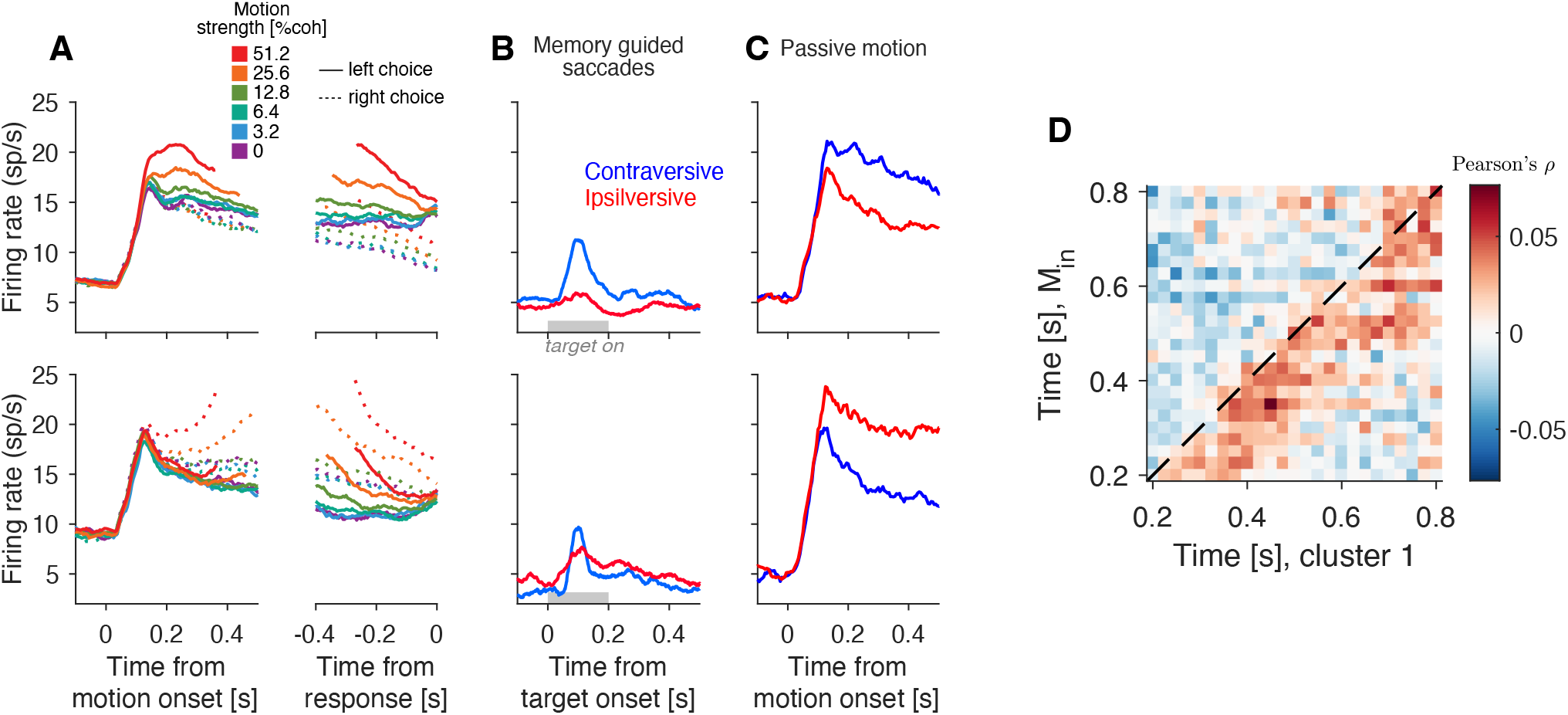
Momentary motion evidence in LIP. **(A–C)** Firing rate of the M_in_ neurons in the random dot motion task (**A**), the memory-guided saccade task (**B**), the passive motion viewing task (**C**). The upper (lower) row represents M_in_ neurons that prefer leftward (rightward, respectively) motion. **(D)** Noise correlations between the motion-selective neurons with the motion stimulus on their response field (ordinate), and the 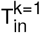 neurons (abscissa). Same conventions as in Fig. 6B–D.

**Figure S8.**
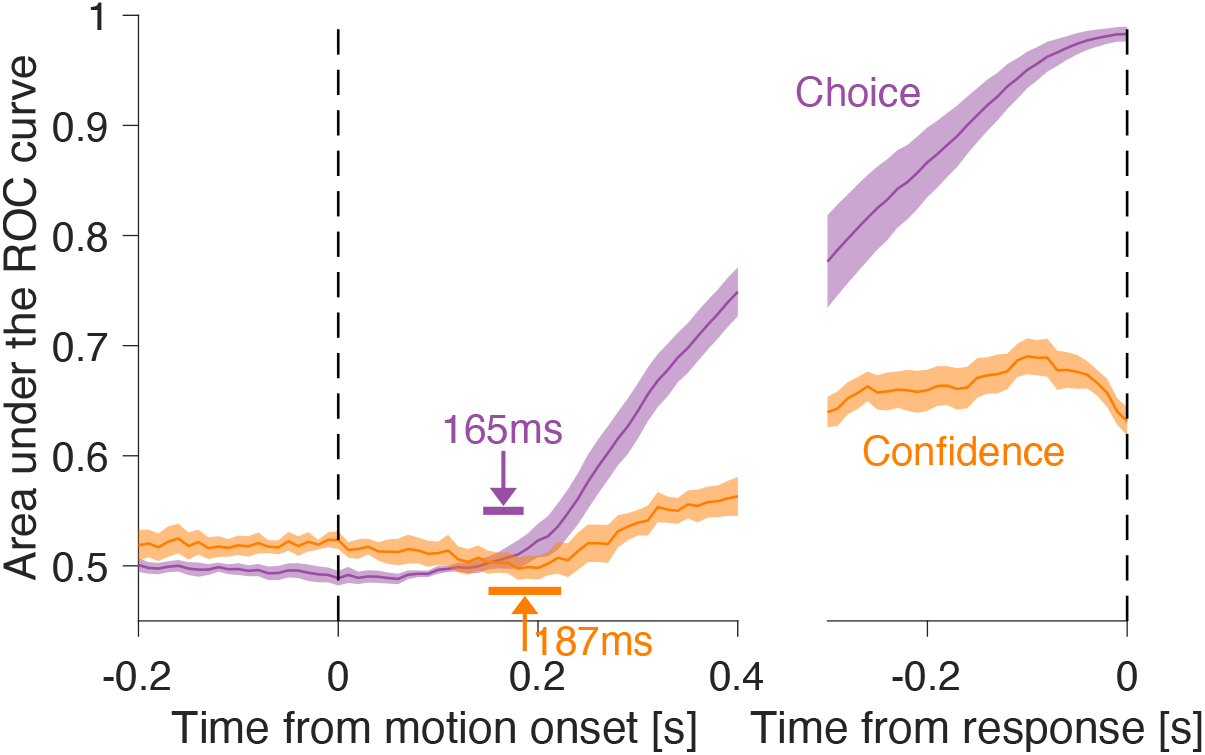
Contemporaneous decoding of choice and accuracy. Time-course of the AUC values obtained from the projection of the neuronal activity along the directions defined by ***β***_**choice**_ (purple) and ***β***_**conf**_ (orange). Shading indicates s.e. across sessions. Projections were calculated in 100 ms windows in steps of 10 ms. The arrows indicate the time when the traces first deviate from baseline (see Methods), and the associated horizontal bars are the s.e. of these estimates.

## Footnotes

1 we use reaction time and choice-response time synonymously, as both refer to the same latency: from onset of the random motion to the onset of the saccadic eye movement used to report the choice. We prefer reaction time to disambiguate behavioral and neural ‘response’ by reserving response for the latter category.

## Notes

**Funding:** Research was supported by the Howard Hughes Medical Institute (M.N.S.) and an R01 grant from the NIH Brain Initiative (M.N.S., R01NS113113).

### Competing Interest Statement

The authors have declared no competing interest.

### Summary of Updates

Changes were made to address reviewers' comments.

